# Proteolysis-free cell migration through crowded environments via mechanical worrying

**DOI:** 10.1101/2020.11.09.372912

**Authors:** Meghan K. Driscoll, Erik S. Welf, Andrew Weems, Etai Sapoznik, Felix Zhou, Vasanth S. Murali, Juan Manuel Garcia-Arcos, Minna Roh-Johnson, Matthieu Piel, Kevin M. Dean, Reto Fiolka, Gaudenz Danuser

## Abstract

Migratory cells often encounter crowded microenvironments through which they must find or make a path. Amoeboid cells are thought to find a path by deforming their bodies to squeeze through tight spaces. Yet many amoeboid cells seem to maintain a near spherical morphology as they move. To examine this unexplored mechanism of migration, we visualized amoeboid melanoma cells in dense environments and found that they carve a path via bleb-driven mechanical degradation of extracellular matrix components without proteolytic degradation. Interactions between adhesions and collagen at the cell front induce a signaling cascade that promotes bleb enlargement via branched actin polymerization. Large blebs abrade collagen, creating feedback between extracellular matrix structure, cell morphology and polarization that enables both path generation and persistent movement.

## Introduction

Cell migration is critical to processes ranging from embryogenesis and wound healing to cancer metastasis.(*1*) When spatially confined, animal and non-animal cells alike exhibit bleb-based motility, a type of amoeboid migration characterized by weak adhesion and minimal proteolytic destruction of the surrounding matrix.(*2, 3*) Amoeboid cells can migrate through tight spaces by deforming their body and nucleus, even to the point of nucleus rupture.(*4–6*) However, many studies of invasive cancer cells, particularly of melanoma, have reported an enrichment of rounded amoeboid cells at the tumor edge.(*7*) Likewise, these cells have been shown to maintain their rounded morphology as they migrate through Matrigel and in tumor xenografts.(*8–10*) Indeed, the pro-migratory effect of elevated intracellular contractility, which is associated with rounding and surface blebbing, is well known.(*7, 10–14*)

Round, amoeboid cells have been observed to move on track-like structures that are sometimes referred to as single-cell invasion tunnels.(*15*) Although under some conditions amoeboid cells secrete matrix metalloproteinases (MMPs),(*16*) amoeboid migration is not usually considered a proteolytic migration mode. Thus, it has been demonstrated that these tunnels can be pre-formed by ‘leader’ cells moving in a mesenchymal manner, a migration mode in which the extracellular matrix (ECM) is degraded via MMPs.(*15*) Despite the notion that amoeboid cells cannot remodel their environment to generate their own paths and thus would be forced to immobility in a very dense microenvironment, blebbing cells in soft extracellular matrices have been observed to physically manipulate fibers, pushing them out of their way(*17*) and tugging on them with adhesions,(*18*) suggesting the possibility of a purely amoeboid migration mode that is also in some way reliant on matrix remodeling.

### Migrating amoeboid cells can carve a path without the need for extracellular proteolytic degradation

To investigate the mechanism of this novel amoeboid matrix remodeling migration strategy, we chose metastatic melanoma cells as a model for bleb-based migration through soft crowded environments. *In vivo*, melanoma has been shown to metastasize to soft environments such as the brain,(*19*) and melanoma cells have often been found to adopt an amoeboid morphology in such 3D environments.(*8, 9, 20*) Consistent with these observations, we noted that melanoma in the soft environment of the zebrafish hindbrain not only have an amoeboid morphology but bleb extensively (Fig. S1A). To enable more systematic imaging studies at high-resolution, we created dense, yet soft *in vivo* mimetics by encapsulating cells in fibrous Collagen 1 gels with bulk stiffnesses on the order of 1 kPa (see Methods).(*21, 22*) We imaged melanoma cells in 3D with light-sheet microscopy at near isotropic ~350 nm resolution, using specially designed sample chambers that do not interfere with the mechanical properties of the mimetics (Fig. 1A, Movie 1).(*20, 23*) Adsorption of collagen to hard surfaces increases collagen stiffness near the surface, (*24–27*) which shifts and eventually diminishes the rounded, blebbing morphotype. To observe migration of these cells without mechanical interference, our chambers enabled cell imaging at greater than 1 mm away from any stiff surfaces.

**Figure 1.**
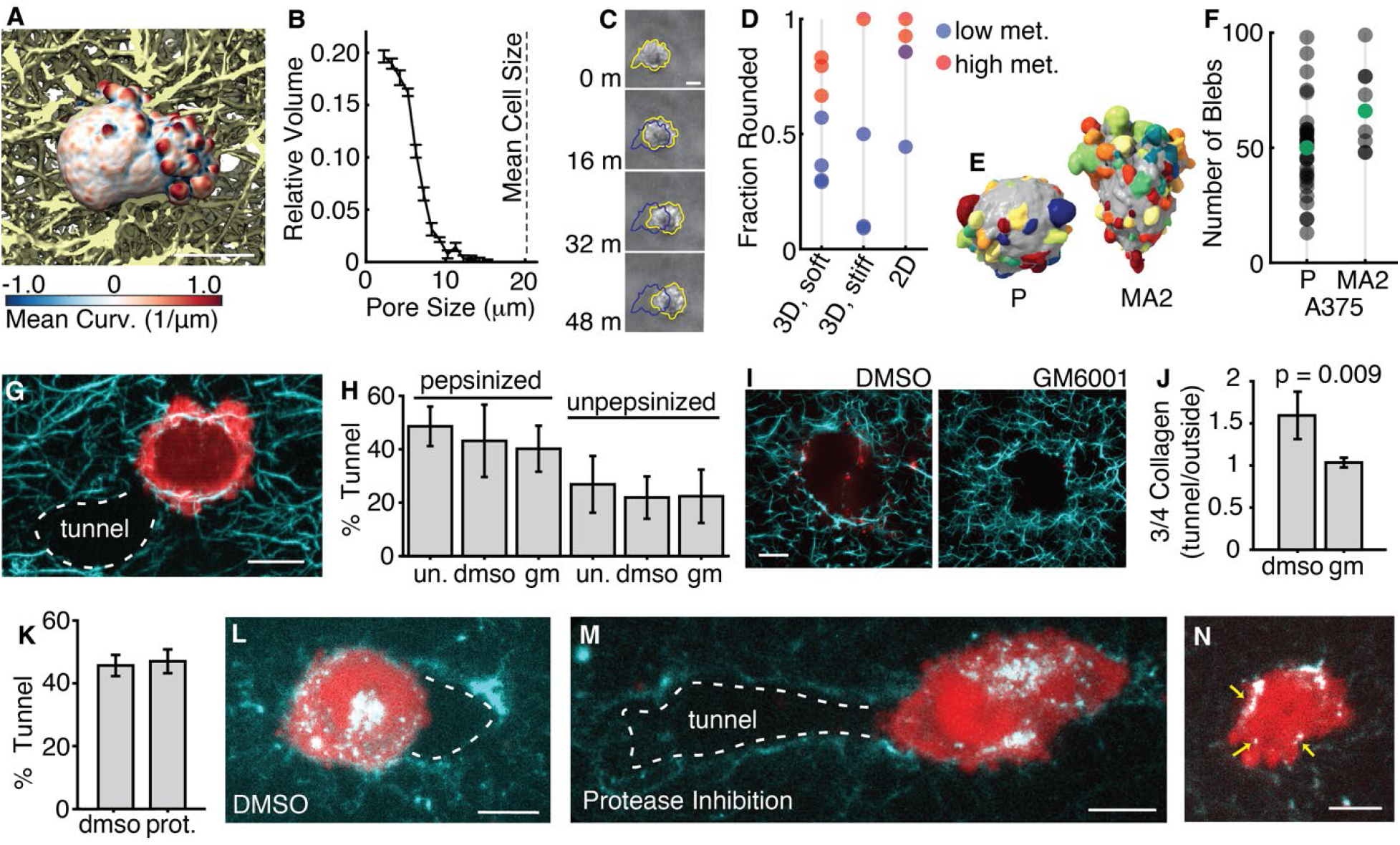
Cells can generate a path through dense collagen without matrix metalloproteinases. (A) Surface rendering of a 3D light-sheet microscopy image of a melanoma cell in 3D collagen with collagen fibers behind it shown yellow. (B) Pore size analysis of 2.0 mg/ml bovine collagen gels showing the fraction of space occupied by a specific pore size (n = 6 gels). Error bars indicate the standard error of the mean. In comparison, the mean diameter of blebby melanoma cells in these types of gels is 20 ± 1 μm (n = 44 cells). (C) Time-lapse phase-contrast microscopy images showing the movement of a round, blebby melanoma cell in 3D collagen. (D) Fraction of melanoma cells with rounded morphology in cell populations extracted from patient-derived xenografts of high vs low metastatic efficiency under three different microenvironmental conditions. (E) Surface renderings of parental, P, and metastatic, MA2, A375 melanoma cells. Computationally identified blebs are shown randomly colored. (F) Number of blebs per cell in the parental, P (n = 33 cells), and metastatic, MA2 (n = 9 cells), cells. Cells with the median number of blebs are indicated in green and shown in panel E. (G) Maximum intensity projection across 4.8 μm of a light-sheet microscope image of an A375MA2 melanoma cell expressing GFP-F-tractin (red) in 3D collagen processed with a steerable filter to enhance edges (cyan). The white dashed line indicates the location of a tunnel. (H) Percent of cells in tunnels after 18-24 hours in pepsinized or unpepsinized bovine collagen. Samples were either untreated (un.; n = 176 cells in pepsinized collagen, 67 cells in unpepsinized collagen), or treated with DSMO (n = 52 cells in pepsinized collagen, 105 cells in unpepsinized collagen), or 40 mM GM6001 (gm; n = 82 cells in pepsinized collagen, 67 cells in unpepsinized collagen). Error bars indicate 95% confidence intervals calculated using the Normal approximation for the confidence interval of a binomial distribution. (I) Single optical sections of 3D Iight-sheet images of collagen samples containing MV3 cells treated with either DMSO or 40 μM GM6001 for ~24 hours. The cyan channel shows collagen with fibers enhanced by a steerable line filter and the red channel shows ¾ collagen antibody fluorescence near tunnels formed by cells. (J) Quantification of ¾ collagen antibody intensity inside a tunnel divided by the intensity outside the tunnel in samples treated with either DMSO or 40 μm GM6001. Error bars show 95% confidence intervals (n = 4 tunnels for each condition). (K) Percent of cells in tunnels after 18-24 hours in pepsinized collagen with cells treated with either DMSO or a protease inhibitor cocktail. (n = 90 cells treated with DMSO and n = 81 cells treated with the protease inhibitor cocktail) (L) An MV3 cell, shown red, in DQ collagen, shown green, treated with DMSO. The DQ collagen appears bright where collagen fibers are coming apart. Maximum intensity projection over 2 μm, which is 12 slices. The white dashed line illustrates the presence of a tunnel behind the cell. (n=11 cells) (M) An MV3 cell, shown red, in DQ collagen, shown green, treated with a protease inhibitor cocktail. Maximum intensity projection over 2 μm, which is 12 slices. (n=20 cells) (N) An MV3 cell, shown red, in DQ collagen, shown green, treated with DMSO. Both images are single slices. Yellow arrows indicate patches of fragmented collagen that are near blebby regions of the cell surface. All scale bars show 10 μm.

To determine whether our *in vivo* mimetics prohibited deformation-based migration through pores, we visualized collagen by sparse fluorescent labeling and measured collagen pore size.(*20, 28, 29*) Although blebs were small enough to fit inside the pores in the collagen network, the nucleus and cell body of amoeboid melanoma cells were too large to fit through the existing pores (Fig. 1A,B, Movie 2), thus rendering a deformation-based mode of migration unlikely. Long-term time-lapse imaging of melanoma cells confirmed that amoeboid cells were nevertheless able to move through the mimetics while maintaining their largely spherical shape (Fig. 1C, Fig. S1B,C).

We next tested whether the amoeboid morphology was associated with melanoma metastatic potential within our *in vivo* mimetic. We imaged populations of primary melanoma cells that were harvested from patients and then passaged in a mouse xenotransplantation system.(*30, 31*) Comparing three mechanically distinct collagen microenvironments, including the mimetic, we found that samples with higher metastatic efficiency were enriched in the amoeboid morphology compared to the stretched mesenchymal morphology (Figure 1D, Figure S1D,E). Furthermore, using unbiased cell shape motif detection,(*32*) we discovered that a parental melanoma cell line (A375P) had a lower average bleb count than a subpopulation of the cell line that had been enriched for metastatic potential in mouse xenografts (A375M2) (Fig. 1E,F).(*33*) Together, these results establish the significance of our experimental system as a model of metastatic cell migration in soft, ECM-dense tissues.

To determine how blebbing melanoma cells migrate through soft, dense collagen, we imaged them 24 hours after seeding. Over this time frame, many cells had created tunnels (Fig. 1G). We observed a similar tunneling phenomenon with multiple melanoma cell lines (Fig. S2A), as well as with pediatric Ewing sarcoma cells (Fig. S2B). Tunnel creation in dense matrices is usually ascribed to matrix metalloproteinase (MMP)-dependent mesenchymal migration.(*34, 35*) Thus, our finding of an amoeboid cell morphology associated with a clearly demarcated, cell-generated path seemed paradoxical in view of the current paradigms of cancer cell motility. To begin to solve this puzzle, we applied a broad spectrum MMP inhibitor, GM6001,(*36, 37*) and found no effect on the ability of melanoma cells to tunnel (Fig. 1H). We confirmed that MMPs act on collagen in proximity to MV3 cells, and that GM6001 inhibited this activity, by measuring the abundance of an antibody that recognizes the MMP-cleaved collagen site in tunnels (Fig. 1I,J) and by direct measurement of MMP enzymatic activity on a synthetic substrate (Fig. S2C). The MMP-independence of the tunneling also held when collagen was not first solubilized using pepsin, which removes a collagen crosslinking site potentially rendering the collagen easier to proteolyze (Fig. 1H).(*38*) Although tunneling is somewhat less frequent in unpepsinized collagen, the pore size is greater as is the extent of directional collagen realignment by the cells, suggesting a reduced need to tunnel (Fig. S3). To test whether a range of proteases beyond MMPs might contribute to tunneling, we used a protease inhibitor cocktail that was previously found to impede migration through tight pores, presumably by blocking proteolytic driven tunneling.(*34*) We again found that inhibiting proteolytic remodeling had no significant effect on tunneling (Fig. 1K). However, as expected, the protease inhibitor cocktail suppressed collagen degradation in the bulk (Fig. S2D). Normalizing for the slightly decreased cell viability (<5%) of the cocktail treated sample compared to the control (Fig. S2E), the cocktail reduced collagen degradation by 86 ± 10% (standard error of the mean). Using high-resolution light-sheet microscopy and a fluorescence marker of collagen fragmentation, DQ collagen, we confirmed that the collagen degradation pattern proximal to the cell appeared to be the same with or without protease inhibition (Fig. 1L,M). Altogether, we found that the local ECM remodeling that enables cell tunneling need not be facilitated by proteolytic degradation of collagen, even while proteases are simultaneously globally remodeling the ECM network.

### Path generation is mediated by bleb-driven ablation of the extracellular matrix (ECM)

To address path generation mechanisms alternative to protease-activity, we analyzed the interactions of cells with collagen in greater detail. We noted that collagen was often fragmented in proximity to blebs (Fig 1N, arrows). Furthermore, cells inside tunnels were usually highly polarized, with many large blebs at the cell front facing the enclosed, collagen-rich end of the tunnel (Fig. 2A, Movies 3, 4). Measuring the difference in frequencies of protrusive and retractive motion on and off blebs, we found that blebs were on average protrusive and non-blebs retractive (Fig. 2B), suggesting that blebs are responsible for cell protrusion through collagen. We next measured the colocalization of blebs with collagen, finding that collagen was enriched in regions near blebs, but not directly on blebs (Fig. 2C). This is explained by bleb interdigitation into pores in the collagen network, resulting in high collagen fiber density at the base of blebs (Fig. 2D). Then we examined collagen motion, which showed movement of individual collagen fibers at the front of tunneling cells (Fig. 2E,F). Using a 3D optical flow algorithm designed to capture multi-scale motion both near and away from cells (Fig. 2G),(*39*) we compared the collagen speed near blebs with the bleb speed for both protruding and retracting blebs (Fig. 2H). For protrusive blebs, at low bleb speeds collagen speed increased linearly with bleb speed, whereas at high bleb speeds collagen motion plateaued, consistent with bleb interdigitation into collagen pores. For retracting blebs we found that even at high bleb speed, collagen was pulled in concert with the blebs, meaning that retracting blebs pull collagen towards the cell surface. Indeed, at the fronts of highly polarized cells, collagen was often enriched into a shell at the cell periphery alongside extensive internalization of labeled collagen (Fig. 2I,J). Over long periods of time, cells slowly agitated the collagen shell, breaking off fragments of the collagen and pulling them into the cell (Fig. 2K).

**Figure 2.**
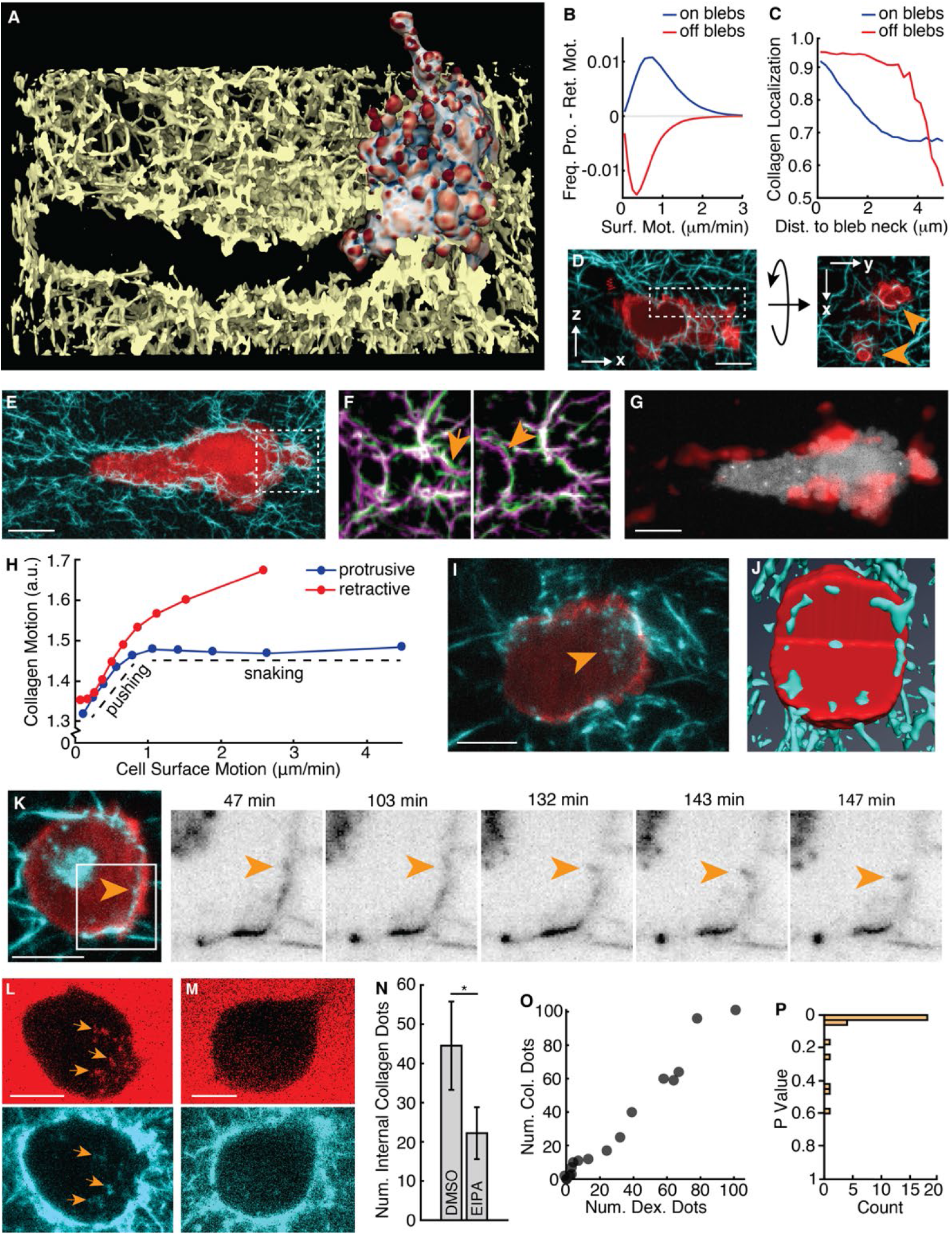
Persistent agitation and internalization of collagen enables cells to dig tunnels. (A) Surface rendering of a light-sheet microscope image of an MV3 melanoma cell, colored by surface curvature, in 3D collagen. The scale for surface curvature is shown in Figure 1A. (B) Frequency of protrusive motion minus the frequency of retractive motion on and off blebs (aggregated over n = 9 cells). (C) Collagen intensity on and off blebs as a function of distance to the nearest bleb neck (aggregated over n= 23 cells). (D) XZ and XY views of maximum intensity projections of light-sheet images of a melanoma cell expressing GFP-AktPH (red) interacting with collagen fibers enhanced by steerable line filtering (cyan) and projected over 5.6 μm and 2.4 μm, respectively. The region shown in the XY view is indicated by the dashed box in the XZ view. Orange arrows indicate blebs interdigitating between collagen fibers. (E) Maximum intensity projection across 3.2 μm of a light-sheet image of a melanoma cell expressing GFP-AktPH (red) in 3D collagen enhanced by steerable line filtering (cyan). The dashed box indicates the region magnified in panel F. (F) Overlay of two different time points (green and magenta, separated by 120 seconds) of a maximum intensity projection over 3.2 μm of a light-sheet microscope image of collagen fibers enhanced by steerable line filtering. Orange arrows indicate the motion of individual collagen fibers. (G) Collagen motion (red), as measured by 3D optical flow, near a melanoma cell expressing GFP-AktPH (white), imaged using light-sheet microscopy and shown as a maximum intensity projection over the entire cell. (H) Collagen motion near the cell surface of blebs associated with either protrusive (blue) or retractive (red) motion (aggregated over n = 5 cells). (I) Maximum intensity projection across 3.2 μm of a light-sheet image of a cell expressing GFP-AktPH (red) in 3D collagen (cyan). The orange arrow indicates internalized collagen at the front of the cell. (J) Surface rendering of a light-sheet microscopy image of a melanoma cell expressing GFP-AktPH (red) in 3D collagen (cyan). A quadrant of the cell and collagen is cut away to show the collagen internalized at the cell periphery. (K) Maximum intensity projection across 3.2 μm of a light-sheet microscope image of a cell expressing GFP-AktPH (red) in 3D collagen (cyan). The dashed box indicates the region magnified in the time-lapse panels to the right and the orange arrow indicates a piece of collagen that is broken off and brought in towards the center of the cell. (L#M) Maximum intensity projection across 3.2 μm of a light-sheet microscope image of MV3 cells (unlabeled) in 3D collagen (cyan) treated with 70 kDa FITC-dextran (red), as well as either DMSO (L) or 50 mM EIPA (M). The orange arrows indicates internalized collagen and dextran in intracellular vesicles. (N) Number of internalized collagen fragments in either DMSO (n = 20 cells) or EIPA-treated (n = 23 cells) cells (difference between conditions per one-sided t-test, p = 0.04). (O) Number of internalized collagen fragments per cell vs. the number of internalized dextran dots in EIPA-treated cells. (P) The p value, calculated for each cell, corresponding to the likelihood that collagen fluorescence intensity is not elevated at the location of dextran dots. All scale bars show 10 μm.

To determine the mechanism of collagen internalization, we incubated cells with high molecular weight dextran, finding that it was ingested alongside labeled collagen (Fig. 2L). Internalization of large liquid-phase molecules is indicative of macropinocytosis, since such molecules are excluded from smaller endocytic vesicles.(*40*) Treatment with the sodium hydrogen exchange inhibitor 5-(N-ethyl-N-isopropyl)amiloride (EIPA), an inhibitor of macropinocytosis,(*41*) decreased the number of dextran-labeled vesicles and internalized collagen fragments, with the number of dextran vesicles and collagen fragments highly correlated across cells (Fig. 2M-O). Testing the hypothesis that collagen localization at detected dextran vesicles was random, we found that the distribution of p-values was heavily tilted towards small values, indicating collagen enrichment within dextran vesicles (Fig. 2P). Furthermore, we did not observe intracellular vesicles containing fluorescently-labeled clathrin light chain (CLC) that were associated with internalized collagen (Fig. S4), indicating that clathrin-mediated endocytosis does not contribute to internalization. Thus, we conclude that macropinocytosis is likely the dominant form of collagen internalization in this mode of path generation. Combined with the known roles in nutrient uptake in depleted cancer microenvironments,(*42*) these results highlight the importance of macropinocytosis in metastatic migration.

### Phosphoinositide 3-kinase (PI3K) establishes bleb polarity in feedback with collagen remodeling

For productive path generation, the slow destruction of dense extracellular matrix at the cell front critically depends on persistence factors that promote the highly polarized and continuous formation of large blebs, which abrade and internalize matrix material in a directed fashion. To measure polarization on the 3D cell surface, we used an approximation of a spherical normal distribution, which has fit parameters that intuitively correspond to the direction of the peak and the peak’s inverse width, here termed the polarization magnitude (Fig. 3A). Using these statistics, we first confirmed that the distribution of large blebs was more polarized than the overall bleb distribution (Fig. 3B). Measuring the directional correlation of large bleb polarization and collagen localization, we next found that large blebs were systematically biased towards areas of high collagen density (Fig. 3C). Hypothesizing that adhesions might couple collagen and bleb localization, we found that paxillin-containing adhesion complexes indeed formed (Fig. 3D, Fig. S5) in the direction of the high collagen density at the closed end of the tunnel (Fig. 3E). A canonical cell polarity factor that is organized by nascent adhesions via focal adhesion kinase (FAK) is PI3K.(*43–45*) Similar to the distribution of adhesions, we observed a striking polarization of PI3K near the closed end of the tunnel (Fig. 3F, Movie 5). The role of PI3K in polarizing non-actin-based protrusions is unclear. PI3K was previously shown to regulate blebbing in *Dictyostelium,(46*) yet a noncanonical form of pressure-based fibroblast motility in 3D microenvironments does not seem to require polarized PI3K signaling.(*47*) Moreover, PI3K signaling was more directionally aligned with large blebs than with blebs of all sizes (Fig. 3G), suggesting that PI3K signaling is involved specifically in the polarization of large blebs. Despite their small size, adhesions in the cortical area persisted for several minutes (Fig. 3H), in contrast to the ~1 min lifetime of similarly-sized nascent adhesions formed in cells on a coverslip (Fig. 3I,J).(*48*) The localization and persistence of cortical adhesions at the front of tunneling cells thus may enable the recruitment of PI3K to the cell front.

**Figure 3.**
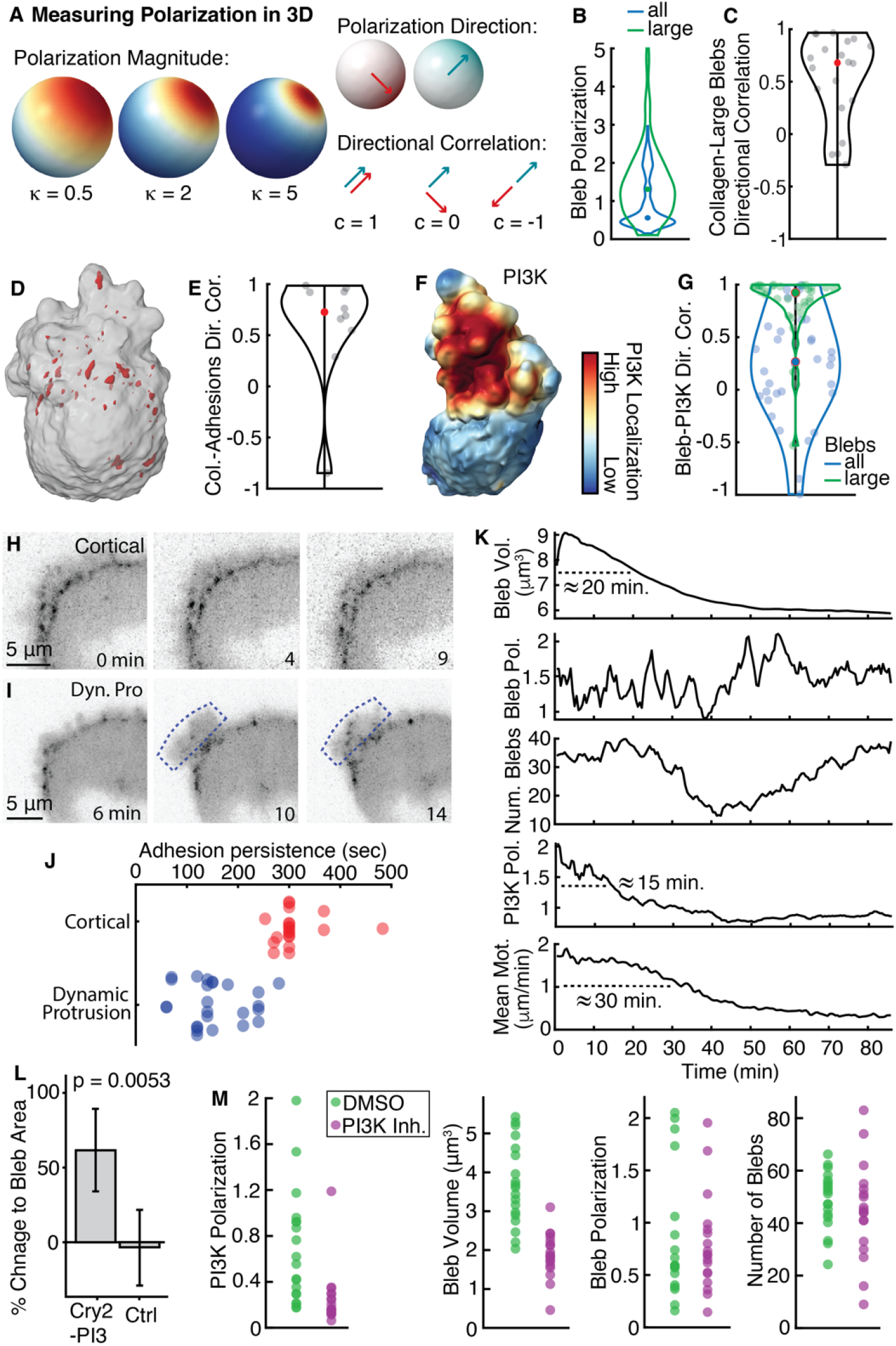
Large blebs are polarized to the cell front via a feedback between collagen remodeling, adhesion formation, and PI3K localization. (A) Simulated species concentrations illustrating example polarization magnitudes and directional correlations. (B) Distributions of polarization magnitudes for all blebs (blue) and for the largest decile of blebs by volume (green). (n = 34 cells) (C) Directional correlation of collagen polarization near the cell surface with polarization of the largest decile of blebs. (D) A 3D surface rendering of a melanoma cell in 3D collagen with the cell surface shown transparent in gray and paxillin-marked adhesions in red. (E) The directional correlation of collagen polarization with adhesion polarization. (F) A surface rendering of a light-sheet microscope image of a melanoma cell in collagen, colored by the localization of GFP-AktPH. (G) The directional correlation of bleb polarization with PI3K polarization for all blebs and for the largest decile of blebs by volume. (H) Time-lapse images of GFP-paxillin adhesions localized to the cortical region in a worrying cell. (I) Time-lapse images of GFP-paxillin adhesions within a dynamic protrusion, indicated by the dashed blue box. (J) Adhesion lifetimes for adhesions near the cell cortex (red) and within dynamic protrusions (blue) in three different cells for each condition. Each data point represents the persistence of a single adhesion. (K) Representative temporal response of blebs and PI3K signaling in an MV3 cell treated with FAK inhibitor 14. From top to bottom, shown are mean bleb volume, bleb polarization magnitude, number of blebs, PI3K polarization magnitude, and mean surface motion magnitude. Dashed lines indicate the approximate full-width half-maximum decay times of measures that are reduced by FAK inhibition. (L) Change in bleb size due to photoactivation, calculated as mean maximum area per bleb during activation divided by mean maximum area per bleb in the same region before activation (p=0.0053, two sided t-test, n=6 regions from 6 cells expressing mCherry-CRY2-iSH2 along with CIBN-CAAX and n=8 regions in 6 wild-type cells). (M) Effect of PI3Kα inhibitor IV compared to DMSO control on bleb and PI3K properties in MV3 cells. PI3K polarization (p = 0.001) and bleb volume (p = 9×10^-8^) show statistically significant differences across treatments, whereas bleb polarization (p = 0.4) and number of blebs (p = 0.12) do not. Statistical testing was performed with a one-sided t-test.

To test this hypothesis, we acutely inhibited FAK signaling using a small molecule inhibitor of FAK-kinase activity. FAK inhibition resulted in a decrease in bleb volume, even though bleb polarization and number were unaffected (Fig. 3K). PI3K polarization and mean cell surface motion were also decreased (Fig. 3K, Fig. S6). Measuring the response times to FAK inhibition, we found that PI3K polarization fell first, followed by bleb volume and then cell surface motion. This led us to conclude that PI3K polarity is upstream of large bleb formation at the cell front. Indeed, stratification of blebs by volume revealed that large blebs, in particular, were enriched for high PI3K signaling and also associated with increased collagen motion (Fig. S7).

To determine if the relationship between PI3K and bleb size was causative, we used photoactivation to increase PI3K signaling locally in blebbing cells, resulting in a striking increase in proximal but not distal bleb size (Fig. 3L, Fig. S8, Movie 6). We also pharmacologically inhibited PI3K signaling by acute addition of a low dose of an inhibitor specific for PI3Kα. In the region of former high PI3K activity, PI3K biosensor intensity and bleb size rapidly decreased, even though de novo bleb formation was not inhibited (Fig. S9, Movie 7). Aggregating over multiple cells, we found that both PI3K polarization and bleb size were decreased by PI3K inhibition, whereas the number of blebs and bleb polarization were not affected (Fig. 3M). Altogether, these results indicate that PI3K is responsible for generating large blebs but does not govern the frequency or location of bleb initiation.

### PI3K enlarges blebs by enhancing branched actin polymerization

The fact that PI3K activity was responsive to FAK manipulation left open the possibility of a reinforcing, mechanical feedback from large blebs to PI3K, which would also explain the observed persistence in polarization of blebs and signaling. To test if the presence of large blebs affected PI3K activity, we reduced cell blebbing via addition of wheat germ agglutinin (WGA), which binds to sialic acid and N-acetylglucosaminyl residues on the extracellular surface of the cell membrane, thereby increasing membrane stiffness.(*49, 50*) We found that decreasing bleb abundance via WGA decreased the polarity and strength of PI3K signaling (Fig. 4A, Fig. S10), supporting the notion of a sustained feedback between blebbing and local PI3K activity.

**Figure 4.**
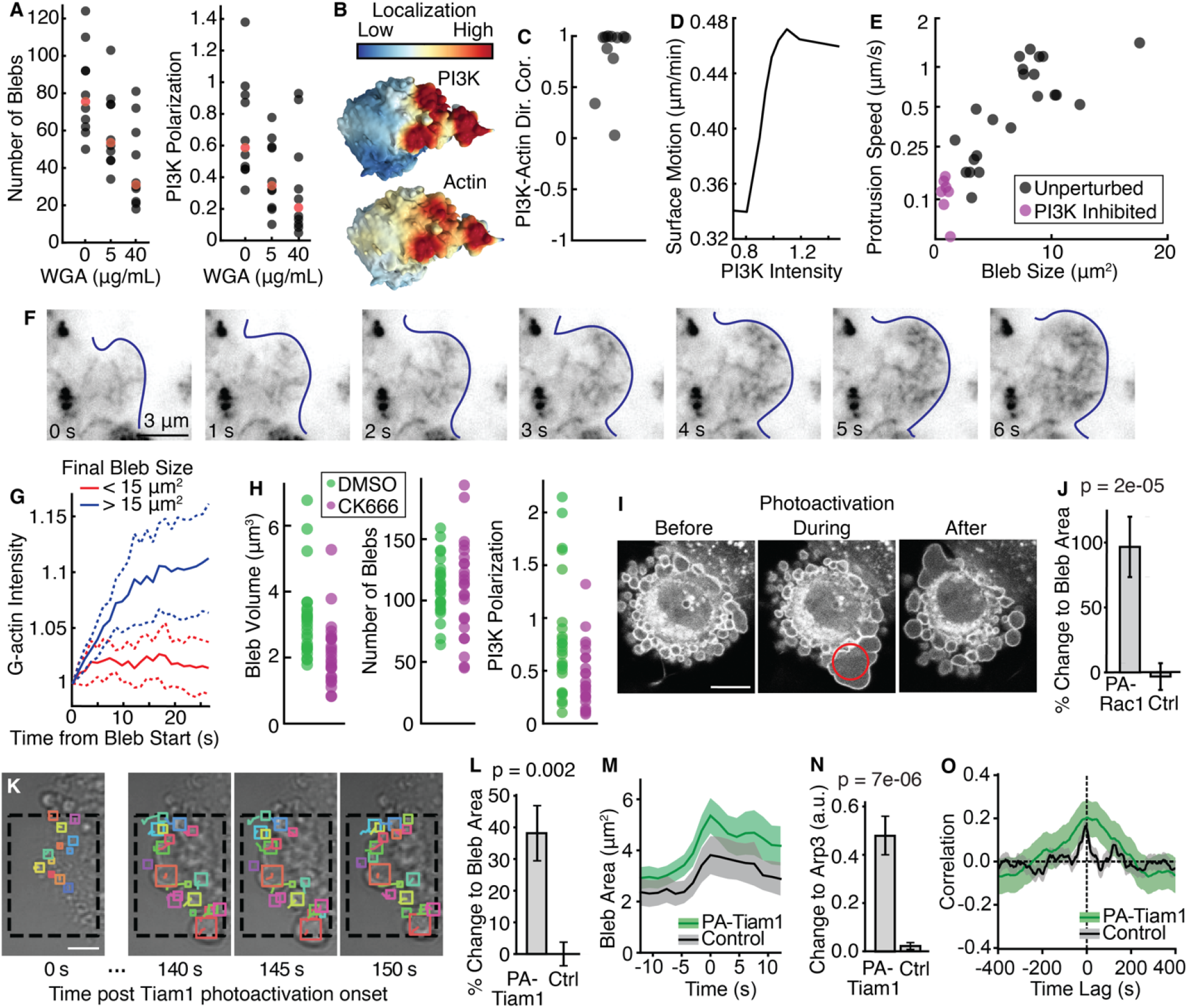
Bleb size is locally controlled by branched actin polymerization. (A) The number of blebs and the PI3K polarization in individual cells as a function of Wheat Germ Agglutinin (WGA) dose. In each category, the value of the median cell is colored red. (B) Surface renderings of 3D light-sheet microscopy images showing GFP-AktPH and HALO-actin intensity on the surface of an MV3 cell. (C) Directional correlation of PI3K and actin polarization in individual MV3 cells (n= 11 cells). (D) Local cell surface motion as a function of local PI3K intensity at the cell surface (n= 9 cells). (E) Protrusion speed of individual blebs as a function of final bleb size in either unperturbed or PI3K inhibited cells. (F) Time-lapse sequence of TIRF microscopy images of a single bleb in a HeLa cell expressing GFP-actin. The bleb edge is indicated by the dark blue line. (G) Mean HALO-actin intensity within large and small blebs in a light-sheet image of MV3 cells as a function of time after bleb initiation. Dashed lines represent 95% confidence intervals (n= 12 blebs from 3 different cells). (H) Effect of Arp2/3 inhibition via CK666 on PI3K and bleb properties, compared to DMSO control. PI3K polarization (p = 0.004) and bleb volume (p = 0.001) show statistically significant differences across samples, whereas the number of blebs do not (p = 0.48). Statistical testing was performed with a one-sided t-test. (I) Spinning disk confocal microscope images showing a single optical slice of mCherry-PA-Rac1 in a mouse embryonic fibroblast before, during and after photoactivation of Rac1 in the area indicated by a red circle. (J) Change in bleb size due to photoactivation, calculated as the mean of the maximum bleb area during activation divided by the mean of the maximum bleb area in the same region before activation (p=1.6 x10^-5^, two-sided t-test; n=6 regions expressing mCherry-PA-Rac1, n=11 regions in cells not expressing mCherry-PA-Rac1). (K) Photoactivation of Rac1 signaling in MV3 melanoma cells using a sspb tagRFP Tiam1 construct. Activation starts at t=0 seconds within the box outlined by the black dashed. Automatically tracked blebs with each bleb shown randomly colored. (L) The change to bleb area for photoactivated regions compared to photoactivated regions in control MV3 cells expressing sspb tagRFP. Error bars indicate the standard error of the mean. (For PA-Tiam1, n=21 regions from 12 cells. For the control, n=19 regions from 7 cells) (M) For individually tracked blebs, the mean bleb area over time with the shaded area indicating the 95% confidence interval. Individual tracks were temporally aligned with maximum bleb area set to t=0. Shown are blebs in photoactivated regions of MV3 cells expressing sspb tagRFP Tiam1 and, as a control, blebs in the same regions prior to photoactivation. (n=416 total blebs extracted from 21 regions and 12 cells.) (N) The change in Arp3 florescence intensity in photoactivated regions of MV3 cells expressing sspb tagRFP Tiam1 compared to photoactivated regions in control cells expressing sspb tagRFP. Error bars indicate the standard error of the mean. (For PA-Tiam1, n=21 regions from 12 cells. For the control, n=19 regions from 7 cells). (O) The cross-correlation of bleb area with Arp3 fluorescence intensity for photoactivated regions of MV3 cells expressing sspb tagRFP Tiam1 compared to photoactivated regions in control cells expressing sspb tagRFP. Shaded areas indicate the standard deviation. (For PA-Tiam1, n=21 regions from 12 cells. For the control, n=19 regions from 7 cells). Unless otherwise indicated, all scale bars are 10 μm.

We therefore sought to uncover the mechanism that couples PI3K activity and bleb size. PI3K is usually associated with actin-driven protrusion.(*51*) In contrast, bleb growth is thought to be driven by pressure-based cytoplasmic flows(*52*) with actin filament formation only at very late growth stages to reform the actin cortex.(*53*) Thus, the involvement of PI3K in bleb expansion seemed paradoxical. Recently, bleb initiation was found to be polarized by actin in zebrafish primordial germ cells, yet in melanoma we found no association of PI3K with bleb number or location, only with bleb size (*54*). We noted, however, that PI3K and actin were directionally correlated (Fig. 4B,C) and that regions of the cell with higher PI3K activity protruded faster (Fig. 4D). Indeed, we found that blebs that ultimately reached a larger size did so by growing faster (Fig. 4E). Inhibition of PI3K activity dramatically decreased the growth rate and final bleb size (Fig. 4E), confirming that fast bleb growth and large size are due to PI3K signaling.

Given the known association of PI3K with actin-based migration, we wondered if actin polymerization could be playing a role in bleb growth despite previous reports otherwise. We found that the F-tractin construct, which localizes to filamentous actin,(*55*) was absent during bleb expansion and localized only to the bleb cortex, as previously reported.(*53*) However, expressing low levels of Halo-tagged actin showed that actin is present during bleb expansion (Fig. S11A). Calculating the ratio of actin to F-tractin revealed that actin was enriched in blebs relative to filamentous actin (Fig. S11B), raising the possibility that nascent actin filaments formed in blebs, but are not recognized by the F-tractin probe. To further explore this possibility, we used a TIRF microscope to image actin in the growing blebs of HeLa cells gene-edited to express a copy of the beta actin gene fused to GFP. We chose this cell model because it allows induction of extended bleb formation by application of slight pressure to the sample chamber. Under these advantageous conditions for high-resolution imaging of the bleb expansion process, we found indeed that actin filaments assemble during bleb growth (Fig. 4F, Movie 8). These data are corroborated in MV3 melanoma cells, where stratification of blebs by their final size indicated a significantly higher concentration of actin in large blebs compared to small blebs, in particular later in their life (Fig. 4G).

The putative link between PI3K signaling, actin polymerization and bleb expansion may rest on the Rac1 – WAVE – Arp2/3 pathway,(*43, 56*) which promotes lamellipodia expansion. To test the involvement of this pathway also in bleb expansion, we first blocked Arp2/3 activity by the small molecule inhibitor CK666,(*57, 58*) which decreased bleb volume but did not alter bleb number (Fig. 4H). This is consistent with previous findings that both genetic and acute inhibition of the Arp2/3 complex reduces bleb size.(*59*) Coupled with previous findings that CK666 treatment does not decrease intracellular pressure(*60*) and thus is not expected to globally decrease the driving force for bleb expansion, our results suggest that active actin polymerization contributes to bleb growth, especially in large blebs. To test this even more directly, and without the risks of altering other Arp2/3-mediated processes, we sought to control bleb growth by acute and local stimulation of the Rac1 signaling pathway using a photoactivatable Rac1 construct (PA-Rac).(*61*) In MV3 cells activation of Rac1 via this probe did not yield reproducible results, likely because the LOV-domain controlled uncaging did not generate sufficiently localized Rac1 activity. We therefore resorted to using mouse embryonic fibroblasts (MEFs) where the functioning of the PA-Rac probe was first demonstrated.(*61*) We noted that a sizable fraction of these MEFs displayed a blebby baseline morphology, similar to amoeboid cell types. In support of our hypothesis that direct stimulation of the Rac1 – WAVE – Arp2/3 pathway would promote bleb expansion, we found that we could systematically and reversibly control bleb volume by photoactivation of Rac1 (Fig. 4I,J). Returning to MV3 melanoma cells, we wondered whether we would be able to enhance the localization of photo-stimulated Rac1 activity by employing a newer generation photoactivatable Tiam1 construct that combines uncaging and sequestration of the active signal.(*62*) Tiam1 is a Rac1-specific guanine nucleotide exchange factor. Indeed, local photoactivation of Tiam1 resulted in a reproducible increase in bleb size that was immediately reversible upon light cessation. To carefully quantify the effect of Tiam1 photoactivation, we developed custom software to track and measure the size of individual blebs (Fig 4K and Movie 7). As with MEFs, these analyses showed that photoactivation of Tiam1 greatly increased bleb size (Fig. 4L). Indeed, measuring bleb area over time for individual blebs, we found that Tiam1 photoactivation increased bleb size, but not duration (Fig. 4M), consistent with our previous findings. In support of our hypothesis that bleb expansion is driven by Rac1 – WAVE – Arp2/3 activation, we found that Tiam1 photoactivation also raises Arp2/3 density in the bleb volume (Fig 4N). To further test the coupling between Arp2/3 recruitment and bleb expansion we computed the temporal cross-correlation between bleb area and Arp2/3 fluorescence in individual blebs. For blebs expanding under Tiam1 activation we found a significant correlation with lag equal 0 seconds, i.e., bleb expansion and Arp2/3 are concurrent (Fig. 4O). Intriguingly, even for the smaller blebs that form without Tiam1 activation the peak cross-correlation, albeit lower, was still significant. This confirms that an Arp2/3-mediated expansion is at play also without experimental manipulation of the Rac1 – WAVE – Arp2/3 pathway. In summary, these results identify that the mechanism by which PI3K localization promotes bleb growth at the front of tunneling cells is promoted by Rac1-mediated actin polymerization.

## Discussion

Our data uncover a mode of cell migration that is effective in dense, soft environments. We refer to this mode as worrying, which means wearing down or tearing repeatedly, like a dog worrying a bone, a meaning that predates the more figurative use as a term related to anxiety.(*63*) In the context of cell migration, worrying describes the mechanism by which sustained agitation and tearing of the extracellular matrix at the cell front by persistently polarized and dynamic cell surface blebs frees a path for the cell to move through a crowded environment. The requirement for persistence is met by a mechanochemical feedback between actin-enforced large bleb formation, matrix ablation, adhesion signaling, and PI3K/Rac1 triggered activation of actin filament assembly inside the bleb. The discovery of this self-reinforcing machinery depended on the development of 3D imaging assays to capture cell dynamic behaviors without mechanical interference from the microscope optics, computer vision to extract the relations between cell blebbing and signaling, and optogenetic approaches to acutely intervene with the feedback loop.

Our custom-built technology enabled the study of migration in dense, yet soft environments. Not only are such tissues common *in vivo*, with melanoma known to prefer soft environments in particular,(*64*) even tissues with higher bulk stiffness are mechanical composites with pockets of mechanically soft microenvironments at the scale of single cells.(*65, 66*) It is well established that mesenchymal migration, which is especially common in stiff environments, is facilitated by ECM remodeling leading to tunnel generation and directional fiber alignment.(*67*) From the recent discovery that amoeboid immune cells build specialized actin structures to push fibers out of the way,(*68*) we now conclude based on our data that all major modes of migration are remodeling modes, in which the environment is at least transiently reorganized. Moving forward, the use of advanced technologies to dissect migration in ever more complex environments will be critical to enhancing our understanding of the mechanisms of environmental remodeling and its integration with other cell biological processes.

## Methods

### Cell culture

MV3 cells were obtained from Peter Friedl (MD Anderson Cancer Center, Houston TX). A375 (ATCC® CRL-1619) and A375MA2 (ATCC® CRL-3223) cells were acquired from ATCC. SKNMC Ewing sarcoma cells were obtained from the Whitehurst lab at UT Southwestern. MV3, A375, and SKNMC cells were cultured in DMEM (Gibco) supplemented with 10% fetal bovine serum (FBS; ThermoFisher) at 37°C and 5% CO_2_.

Populations of primary melanoma cells were created from tumors grown in murine xenograft models as described previously (*30*), and provided as a gift by the Laboratory of Sean Morrison (UT Southwestern Medical Center, Dallas, TX). Briefly, cells were suspended in Leibovitz’s L-15 Medium (ThermoFisher) containing mg/ml bovine serum albumin, 1% penicillin/streptomycin, 10 mM HEPES and 25% high protein Matrigel (product 354248; BD Biosciences). Subcutaneous injections of human melanoma cells were performed in the flank of NOD.CB17-Prkdcscid Il2rgtm1Wjl/SzJ (NSG) mice (Jackson Laboratory). These experiments were performed according to protocols approved by the animal use committees at the University of Texas Southwestern Medical Center (protocol 2011-0118). After surgical removal, tumors were mechanically dissociated and subjected to enzymatic digestion for 20 min with 200 U/ml collagenase IV (Worthington), 5 mM CaCl_2_, and 50 U/ml DNase at 37°C. Cells were filtered through a 40 μm cell strainer to break up cell aggregates and washed through the strainer to remove cells from large tissue pieces. The cells were then cultured in medium containing the Melanocyte Growth Kit (ATCC PCS-200-042) and Dermal Cell Basal Medium (ATCC PCS-200-030).

### Inhibitors and other reagents

GM6001 was purchased from Sigma (CC1010). The ¾-collagen antibody was purchased from Adipogen (AG-25T-011). The components for the protease inhibitor cocktail, which was previously detailed by Wolf *et. al*. (*34*), were purchased from Selleckchem and included BB-2516 (S7156), a matrix metalloprotease (MMP) inhibitor for MMP-9, MMP-1, MMP-2, MMP-14 and MMP-7; E-64 (S7379) an irreversible and selective cysteine protease inhibitor that also inhibits papain, cathepsins B and H, and calapin; Pepstatin A (S7381) and aspartic protease inhibitor that also inhibits cathepsins D and E; Leupeptin (S7380) a reversible inhibitor of serine and cysteine proteases; and Aprotinin (S7377) a serine protease inhibitor. DQ Collagen was purchased from Thermo Fisher Scientific (D12060), as was Ethidium Homodimer (E1169). Hoechst 33342 was purchased from Molecular Probes (H3570). FAK inhibitor 14 was purchased from Tocris (3414). PI3K alpha inhibitor IV was purchased from Santa Cruz (sc-222170). CK666 was purchased from Millipore Sigma (SML0006). NEIPA was purchased from Sigma (A3085). WGA was purchased from VWR (101098-084). 70 kDa Fluorescein isothiocyanate–dextran was purchased from Sigma (46945).

### Protease inhibition assays

To test the efficiency of MMP inhibition via treatment with GM6001, we used an MMP activity assay kit by Abcam (ab112146) in accordance with their instructions. MV3 melanoma cells were cultured in a 24 well plate with (a) media with no cells, (b) cells treated with vehicle (DMSO) control, and (c) cells treated with GM6001 for 24h. Two positive controls of recombinant human MMP1 and MMP8 from RnD systems (901-MP & 908-MP) were used for the assay. The control MMPs were dissolved in assay buffer and a 2mM AMPA working solution was prepared with assay buffer. The MMP and the test samples were mixed 1:1 vol/vol with the AMPA working solution and incubated for 1h at 37 °C. The MMP green substrate working solution was prepared in assay buffer and then mixed 1:1 vol/vol in the black walled 96 well plate and further incubated for 1h. The samples were then read on a Biotek, Synergy H1 hybrid plate reader at Ex/Em = 490/525 nm. The effectiveness of GM6001 was visually confirmed via high-resolution light-sheet microscopy using a ¾-collagen antibody that binds to fragmented collagen.

To test the efficiency of the broad spectrum protease inhibitor cocktail described in the previous section, MV3 melanoma cells were cultured in 2 mg/mL bovine collagen with 2% FITC labeled DQ collagen. Immediately upon collagen polymerization, the melanoma cells were treated with the protease inhibitor cocktail or DMSO vehicle control for 48 hours. The protease inhibitor cocktail constituted BB2516 (100 μM), E-64 (250 μM), Pepstatin A (100 μM), Leupeptin (2 μM), and Aprotinin (2.2 μM). The final concentration of DMSO in the cocktail was 0.24%. After 48 hours the cells embedded in the collagen were pelleted and the supernatant was collected. The absorbance of the supernatant was evaluated in the BioTek plate reader with an excitation of 480 nm and an emission of 525 nm. The fluorescence was normalized to the DMSO control.

As a further control, we measured the effect of the protease inhibitor cocktail on cell viability. MV3 melanoma cells were cultured in 3D collagen matrices in 96 well plates as previously described by Murali *et al*. (*69*). Briefly, cells were seeded at 30,000 cells/well in bovine collagen at a final concentration of 2mg/mL. Upon collagen polymerization, the cells were treated with either the vehicle control or the protease inhibitor cocktail for 48 hours. A 3D viability assay was then performed by incubating the cells with ethidium homodimer at a final concentration of 4 μM, labeling dead cells, and Hoechst 33342 at a final concentration of 15 μg/mL, labeling all nuclei, in phenol red free DMEM supplemented with 10% FBS for 30 mins. Imaging was then performed of the cells with a Nikon-Ti epifluorescence microscope with an OKO temperature control and CO_2_ control system regulated at 37°C and 5% CO_2_ as previously described (*69*). The images were quantified using Cell Profiler (*70*) by evaluating the total number of positive pixels for each channel.(*71*) Cell death was quantified as the ratio of ethidium homodimer pixels to Hoechst pixels. We found that for the protease inhibitor cocktail treated cells 16 ± 3% were non-viable, whereas for the vehicle treated cells, 12 ± 3% were non-viable. This small, albeit statistically significant difference (p-value: 0.008, two-sided paired t test), does not explain the large suppression of collagen degradation in the cells treated with protease inhibitor cocktail.

### Recombinant DNA Constructs

The GFP-AktPH construct was obtained from the laboratory of Jason Haugh (North Carolina State University, Raleigh NC)(*72*) and cloned into the pLVX-IRES-puro vector (Clontech). The GFP-tractin construct was a gift from Dyche Mullins (Addgene plasmid # 58473; http://n2t.net/addgene:58473; RRID:Addgene_58473) (*73*) and was cloned into the pLVX-IRES-puro vector (Clontech). Paxillin-pEGFP was a gift from Rick Horwitz (Addgene plasmid # 15233; http://n2t.net/addgene:15233; RRID:Addgene_15233) (*74*). mRuby2-CLC was a gift from the laboratory of Sandra Schmid (UT Southwestern Medical Center). Cells expressing lentiviral vectors were created by following the manufacturer’s instructions for virus preparation and cell infection (Clontech). Cells were selected for expression by treatment with puromycin, G418, or by fluorescence activated cell sorting.

The photoactivatable PI3K construct (Idevall-Hagren et al., 2012) was created by cloning mCherry-CRY2-iSH2 (Addgene Plasmid #66839) into the pLVX-neo vector (Clontech). The CIBN-CAAX plasmid was obtained from Addgene (Plasmid #79574) and cloned into the pLVX-puro vector. Cells expressing both the mCherry-CRY2-iSH2 and the CIBN-CAAX constructs were selected by treatment with 10 mg/mL puromycin and fluorescence activated cell sorting. It is critical for the two-part cry2 photoactivation system that cells express sufficient concentration of the CIBN-CAAX construct or the cry2 construct will aggregate in the cytosol instead of being recruited to the membrane. Thus, the optimal ratio of CIBN:cry2 is greater than one; cells expressing insufficient CIBN-CAAX will not respond to light. We also noted through the course of our experiments that cells will stop expressing one or both of these constructs if not kept constantly under selective pressure. Such a loss of expression will result in non-responsive cells.

Rac1 activation was achieved by expressing PA-Rac1 via retro-virus infection using pBabe-TetCMV-puro-mCherry-PA-Rac1 (Addgene Plasmid #22035) in MV3 cells. After sorting cells for high expression, photo-stimulation of blebby cells yielded no change in cell morphology. Since this construct was originally shown to work in mouse embryonic fibroblasts (MEFs),(*61*) we expressed the same PA-Rac1 construct in MEFs and photo-stimulated a sub-population of blebby cells, which resulted in a dramatic and local increase in bleb size.

To attempt to activate Rac1 in MV3 cells, we decided to test a newer generation photoactivatable construct of Tiam1, a Rac1-specific guanine nucleotide exchange factor.(*62*) The light-induced localization of Tiam1 was achieved by expressing two DNA constructs in MV3 cells, pLL7.0: mTiam1(64-437)-tgRFPt-SSPB WT (Addgene Plasmid #60417) and pLV_Lyn11-mTurquoise2-iLID (Addgene Plasmid #161000). Cells were also infected to express CMV100-Arp3-Halo. To do so, C-terminally Halo-tagged human Arp3(*75*) was PCR amplified with primers: 5’-tatataactagtgccaccatggcgggacggctg −3’ and 5’-tatataacgcgtttaaccggaaatctcgagcgtcg −3’, cloned into the SpeI/MluI sites of pLVXCMV100(*76*) and sequence verified. Before sorting and imaging, cells were labeled with Janelia Fluor® HaloTag® 646 ligand. Cells expressing three components were selected by fluorescence activated cell sorting and their response was assessed by localization of TgRFPt to the cell membrane upon 488 nm local illumination. For control experiments, cells were infected with pLL7.0: tgRFPt-SSPB R73Q (Addgene Plasmid #60416), which provided for the light-induced localization of tgRFPt without Tiam1.

Overexpression of fluorescently tagged monomeric actin can perturb cell cytoskeletal dynamics. To avoid this artifact while imaging tagged actin, we expressed HALO-tagged actin under the control of a truncated CMV promotor, which results in lower expression of tagged actin than the full length promoter. The original actin construct features an 18 amino acid linker between mNeonGreen and actin in a pLVX-shRNA2 vector and was obtained from Allele Biotech. We truncated the CMV promoter, and replaced the mNeonGreen fluorophore with the HALO tag sequence. The sequence of the CMV100 promoter region is as follows, with the CMV sequence underlined and the start codon in bold: <u>AGTTATTAATAGTAATCAATTACGGGGTCATTAGTTCATAGCCCATATATGGAGTTCCGCGTTACATAACTTACGGT AAATGGCCCGCCTGGCTGACCG</u>CCGCTAGCGCTAACTAGT**GCCACCATG**

### Phase-contrast imaging

Live-cell phase-contrast imaging was performed on a Nikon Ti microscope equipped with an environmental chamber held at 37°C and 5% CO_2_ and imaged with 20x magnification.

### Cells on top of gels

Collagen slabs were made from rat tail collagen Type 1 (Corning; 354249) at a final concentration of 3 mg/mL, created by mixing with the appropriate volume of 10x PBS and water and neutralized with 1N NaOH. A total of 200 μL of collagen solution was added to the glass bottom portion of a gamma irradiated 35 mm glass bottom culture dish (MatTek P35G-0-20-C). The dish was then placed in an incubator at 37°C for 15 minutes to allow for polymerization. Cells were seeded on top of the collagen slab at a final cell count of 5000 cells in 400 μL of medium per dish. The dish was then placed in a 37°C incubator for 4 hours. Following incubation, 1 mL of medium was gently added to the dish. The medium was gently stirred to suspend debris and unattached cells. The medium was then drawn off and gently replaced with 2 mL of fresh medium.

### Cells embedded in 3D collagen

Collagen gels were created by mixing bovine collagen I (Advanced Biomatrix 5005 and 5026) with concentrated phosphate buffered saline (PBS) and water for a final concentration of 2 mg/mL collagen. This collagen solution was then brought to pH 7 with 1N NaOH and mixed with cells just prior to incubation at 37°C to induce collagen polymerization. Cells were suspended using trypsin/EDTA (Gibco), centrifuged to remove media, and then mixed with collagen just prior to incubation at 37°C to initiate collagen polymerization. To image collagen fibers, a small amount of collagen was conjugated directly to AlexaFluor 568 dye and mixed with the collagen sample just prior to polymerization. FITC-conjugated collagen was purchased from Sigma (C4361).

### 3D confocal imaging

The cell/collagen mixture described in the previous section was added to Nunc Lab-Tek II Chambered coverglass samples holders with a No. 1.5 borosilicate glass bottom (Thermo Scientific). Cells were fixed with paraformaldehyde and stained with Hoechst and FITC-phalloidin. Images were acquired on a Zeiss LSM 880 using a plan-apochromat 63x/1.4 oil objective.

### Classification of cell morphology

In Figure 1D, we classified cell morphologies as either amoeboid (rounded) or not amoeboid. For cells imaged at high-resolution, either via confocal or light-sheet microscopy, we labeled cells with extensive blebbing, as well as cells that were rounded, as amoeboid. For cells imaged at lower resolution via phase contrast microscopy, we labeled only cells that were rounded as amoeboid, since the resolution was insufficient to reliably visualize blebs.

### Zebrafish injection and imaging

B16F10 melanoma cells expressing Lifeact-eGFP were injected into the hindbrain ventricle of 2 days post-fertilization wildtype zebrafish larvae using previously described protocols.(*77*) Briefly, B16F10 melanoma cells were suspended in HBSS. 25-50 cancer cells were transplanted into the hindbrain ventricle of anesthetized larvae. Injected zebrafish larvae were incubated at 31°C with 0.2 mM PTU to prevent pigment formation. Live-cell *in vivo* imaging was performed using a Zeiss spinning disc microscope with a QuantEM EMCCD camera.

### TIRF microscopy

Human cervical adenocarcinoma cells HeLa-Kyoto with TALEN-edited ActB fused with GFP (Cellectis, France) were maintained in DMEM/F12 supplemented with 10% FBS (Invitrogen) at 37°C and 5% CO2, and imaged using CO2-independent medium (Invitrogen) supplemented with 10% FBS. Cells were confined by a PDMS stamp in a non-adhesive, PLL-g-PEG (0.5mg/ml) coated chamber of ~3⍰m height, as described previously (*78*), and imaged 20 minutes after initiating the confinement to observe actin dynamics within blebs. The high NA TIRF consisted of a standard setup equipped with a 473nm laser 500 mW (Laserquantum), an objective TIRF NA=1.49 (Olympus), and a camera (Andor Zyla 4.2). A single notch filter was used in the emission light path to block the laser line at 473 nm (Chroma). Acquisition was controlled by the Andor SOLIS software.

### 3D cell tracking from phase-contrast movies

Cells were embedded in 2.0 mg/mL pepsinized bovine collagen in Nunc Lab-Tek II Chambered Coverglass samples holders as described above. Live-cell phase-contrast imaging was performed on a Nikon Ti microscope as described above. Cells were outlined manually using ImageJ, and position and shape data were exported for analysis using Matlab. Cell shape was calculated using roundness, given by 4*area/(π*major_axis^2), and cells were classified as either round (roundness > 0.8) or stretched (roundness < 0.8). Autocorrelation was calculated using the Matlab function *xcorr*. Cell velocity was calculated from cell centroid positions.

### Photoactivation

Photoactivation of subcellular regions was performed using a 488 nm laser at 10% power via the FRAP module of a Zeiss LSM780 outfitted with temperature and CO_2_ control. Cells for the PI3K optogenetics experiments were treated with 200 μM of PI3K inhibitor IV just prior to photo activation. To assess bleb size change in phase contrast movies of PI3K photoactivation, we analyzed multiple blebs within the stimulated region by manually outlining individual blebs at their largest size using ImageJ. Bleb size was measured prior to activation and during activation in the same sub-region of the cell. Similarly, PA-Rac1 activation in MEFs was also performed on a Zeiss LSM780 with local FRAP activation and assessed using manual outlining of blebs prior to and after photoactivation.

Cells for Tiam1 photoinduced localization were imaged with a Zeiss LSM780 microscope and analyzed for bleb size via phase contrast and tagRFP-T imaging and for Arp2/3 recruitment via Janelia Fluor® HaloTag® 646. Bleb dynamics was followed 10 minutes prior to and 10 minutes after onset of 488 nm photo-activation with a 5-10 second imaging interval. Bleb and cytoskeletal dynamics were analyzed using a custom-built analysis package as described below.

### 3D high-resolution light-sheet imaging

3D samples were imaged using either an axially-swept light-sheet microscope(*23*) or a meSPIM microscope,(*20*) both of which provide nearly isotropic, diffraction-limited 3D images. The sample holder was specially designed to allow imaging of cells in soft environments and away from hard surfaces (*79*). Samples were imaged in phenol red free DMEM containing 25mM HEPES (ThermoFisher) with 10% FBS and antibiotic-antimycotic (Gibco), held at 37°C during imaging. Images were collected using sCMOS cameras (Orca Flash4.0 v2, Hamamatsu) and microscopes were operated using custom Labview software. All software was developed using a 64-bit version of LabView 2016 equipped with the LabView Run-Time Engine, Vision Development Module, Vision Run-Time Module and all appropriate device drivers, including NI-RIO Drivers (National Instruments). Software communicated with the camera via the DCAM-API for the Active Silicon Firebird frame-grabber and delivered a series of deterministic TTL triggers with a field programmable gate array (PCIe 7852R, National Instruments). These triggers included analog outputs for control of mirror galvanometers, piezoelectric actuators, laser modulation and blanking, camera fire and external trigger. All images were saved in the OME-TIFF format (https://docs.openmicroscopy.org/ome-model/5.6.3/ome-tiff/). Microscope software is available upon completion of a Material Transfer Agreement with the University of Texas Southwestern Medical Center Tech Transfer office.

### 3D cell image analysis

3D light-sheet images of cells were first deconvolved using the Richardson-Lucy algorithm built-in to Matlab (Mathworks). To reduce deconvolution artifacts, images were apodized, as previously described (*20*). Following deconvolution, we used our previously published u-shape3D analysis framework (*32*) to segment cells, detect blebs, map fluorescence intensity to the cell surface, measure surface motion, and calculate polarization statistics. Briefly, images of cells were segmented to create a cell surface represented as a 3D triangle mesh. We used u-shape3D’s *twoLevelSurface* segmentation mode, which combines a blurred image of the cell interior with an automatically thresholded image of the cell surface. Blebs were detected by decomposing the surface into convex patches, and using a machine learning algorithm to classify the patches as a bleb or not a bleb. For each patch classified as a bleb, the bleb neck was defined as the boundary between that patch and neighboring patches. Distance from a bleb neck was calculated at every face on the mesh as the geodesic distance to the closest bleb neck. To determine the fluorescence intensity at each mesh face, we used the raw, non-deconvolved, fluorescence image. At each mesh face, a kd-tree was used to identify the cell-interior voxels within a sampling radius of 1 or 2 μm of the mesh face. Before averaging the intensity values in these voxels, the intensity values were depth-normalized to correct for surface-curvature dependent artifacts (*80*). The u-shape3D software, as well as the trained machine learning models used here, are available with the previously published manuscript (*32*).

Polarization statistics were calculated by mapping data defined on the cell surface to a sphere, and fitting the mapped data to a 3D von Mises distribution, which is akin to a spherical normal distribution. We calculated bleb polarization by representing each bleb by the location on the bleb surface farthest from the bleb neck, with distances measured on the cell surface. Additionally, since the adhesion images had substantial fluorescence background, to measure adhesion polarization, we bandpass-filtered the raw images via a difference of Gaussians procedure, selecting for objects between 1 and 6 pixels in radius.

### 3D collagen image analysis

To enhance linear image features, such as collagen fibers, the 3D collagen images were processed with a steerable filter of width 2 pixels, as previously described (*20*). To emphasize collagen fiber location, some figure panels, as indicated in the figure legends, show steerable-filter enhanced collagen. Other collagen images, especially those related to endocytosis, were neither filtered nor deconvolved to avoid the creation of artifacts. Collagen polarization near the cell surface was measured after mapping image intensity values from steerable-filtered images onto the cell surface. Following steerable filtering and automatic thresholding, the nematic order parameter of collagen networks was calculated as described previously(*20*), except that the average fiber directionality in each 3D image was used as the reference direction. The fiber directionality was calculated at each voxel via a steerable filter. Collagen pore size analysis was also performed as described previously (*29*). Images were filtered and then thresholded at 2.5 times the intensity threshold calculated by Otsu’s algorithm (*81*). To measure pore sizes, for each image, we first fitted the largest possible sphere into the collagen pores. We then iteratively fitted the next largest sphere into the pore space minus the volume of previously fitted spheres until no remaining spheres above a size threshold would fit. We defined the distribution of collagen pore sizes as the distribution of fitted sphere diameters. Collagen motion was measured using a previously published 3D optical flow algorithm (*39*). This algorithm combines a matching framework for large displacements across frames with a variational framework for small displacements. We mapped the magnitude of the collagen motion calculated via optical flow onto the cell surface using the framework for mapping fluorescence intensity onto mesh faces described above. To create panel 2H, we separated cell surface motion, aggregated across multiple cells, into bins by magnitude and found the mean collagen motion magnitude associated with each bin. The collagen sample moves during imaging, and although the average collagen motion in each frame was subtracted from the measured collagen velocities, residual local motions contribute to create a non-zero background collagen motion.

### 3D dextran assay image analysis

To measure the uptake of collagen fragments alongside 70 kDa dextran, we first segmented the cell using the dextran channel. To do so, we inverted each 3D image, subtracted the median intensity, normalized by the 99^th^ intensity percentile, subtracted the image background, thresholded, morphologically dilated by 1 pixel, morphologically eroded by 8 pixels, filled holes, and finally selected the largest image component. Since the cell is morphologically eroded to a greater extent than it is dilated, the cell segmentation is effectively shrunk, reducing the effect of segmentation errors on later analysis. To detect dots of endocytosed collagen and dextran, we employed a previously published multiscale particle detector (*82*). We used scales of 1.5 to 4 pixels, an ⍰⍰= 0.01, and detected dots only inside the segmented cell. The p-value distribution shown in Figure 2O results from testing, for each cell, the hypothesis that collagen fluorescence intensity is greater at the location of detected dextran dots than elsewhere in the cell. To calculate the p value for each cell, we randomly picked *n* collagen intensity values within the cell 100,000 times, where *n* is the number of detected dextran dots, and calculated the probability that the mean of the randomly picked values was greater than the mean of the collagen intensity values at the true detected dextran dots.

### Image analysis of photoactivation

To assess the temporal bleb size change in phase contrast movies before and after local photoactivation, we tracked and analyzed all detected blebs within the stimulated region automatically using a custom Python script. Briefly, video frames were first Gaussian smoothed with = 1 pixel, histogram equalized to enhance edges and intensity inverted if the blebs were darker than the cell. Blebs were subsequently detected as circular features frame-by-frame using a multiscale difference of Gaussian (DoG) blob detection approach (*83*). For a reference pixel resolution of 0.263 m, the detector used a Gaussian pyramid scheme with minimum = 2 pixels, maximum = 12 pixels and ratio of 1.2. For videos collected of different pixel resolution, the DoG detector parameters were manually adjusted on an individual basis, generally by scaling the parameters by a multiplicative scaling factor equal to the current pixel resolution / 0.263 m to ensure maximal bleb detection coverage in the stimulated sub-region. Detected blebs were linked over time into tracks using bipartite matching using the intersection over union of bleb bounding boxes as the distance function between pairs. To handle large changes in size, the matching between the current and next frame was carried out on the bounding box coordinates of the bleb as predicted by local optical flow (*84*). In case of temporary occlusion or missed detection, any non-matched bleb detections were propagated for up to 5 frames (15-25 s) according to estimated optical flow before track termination. Tracks with > *2* frames (10s, comparable to an average bleb lifetime) were retained for analysis. For each individual tracked bleb the 2D bleb area timeseries was measured as with the radius, corresponding to the maximal DoG filter response fired by the bleb signal. Corresponding normalized molecular activity within the bleb was measured as the mean fluorescence intensities within the bleb bounding box divided by the mean fluorescence intensity of all pixels comprising the cell border. To assess the temporal growth profile of an average bleb before and after photoactivation separately for Tiam1 and control constructs, we first temporally aligned all individual bleb tracks pooled across all movies. For each construct condition, individual bleb tracks were sorted as before or after according to whether the timepoint of maximal bleb area occurred before or after photoactivation. Individual track timeseries were then resampled to the same temporal sampling using linear spline interpolation and a variance filter (threshold 0.05) was used to remove non-changing erroneous tracks. Tracks were temporally aligned using the timepoint of maximal bleb area as timepoint 0, taking a window of 8 timepoints on either side (a total 25s). To avoid any extremal values that may bias the average profile, any tracks with a maximum area > mean s.t.d was removed prior to computing the final mean bleb area 95% confidence interval curves in Figure 4M. To test if we see statistically significant changes across individual photoactivated cell sub-regions, we used mean statistics. To statistically assess bleb size change across all sub-regions across multiple cells, the mean largest area of individual tracked blebs were computed per sub-region. For fluorescence signals we computed the mean bleb intensities prior to and after photoactivation. Two-sided paired *t*-test,, was then used to determine significant differences in the ensemble mean area across sub-regions. Given the short median bleb track length of 9 frames (13 seconds), to statistically verify temporal coincidence between i) bleb size change and photoactivation and ii) bleb size change and Arp3 recruitment, we evaluated cross-correlations on the average time series of all blebs falling into the same subregion. The ensemble mean time series across sub-regions were first resampled to the same temporal frequency using linear spline interpolation and the zero-mean normalized cross-correlation curves were computed per sub-region. Sub-region cross-correlation curves were averaged at all time-lags to derive the mean s.d. cross-correlation curve. A deviation of the curve greater than the s.d. interval at a time lag of 0 indicates significant instantaneous correlation.

### Visualization and Statistics

3D surface renderings were made in ChimeraX (*85*). Colored triangle meshes representing the cell surface were imported into ChimeraX from u-shape3D as Collada dae files, as previously described (*32*). To render collagen, steerable-filtered images were opened directly in ChimeraX and thresholded. To create the rendering of adhesions shown in Figure 3D, the raw paxillin images were bandpassed, admitting objects between 0.5 and 3 pixels in radius, and then median filtered.

Figure 2H has histograms with varied bin sizes. To avoid the existence of bins with very little data, each bin in this panel contains a decile of data. Furthermore, to ease visual interpretation, the time series data in Figure 3K were smoothed using a moving average filter with a span of 5 frames.

All statistical comparisons shown in figures were calculated using a one-sided or two-sided t-test with ⍰⍰= 0.05. Error bars in figures show either 95% confidence intervals or the standard error of the mean, as stated in the figure legends. Number of cells and/or number of different experiments analyzed are given in the figure legends.

## Supporting information

Video collection

## Acknowledgments

We thank Allan Zhang for manually tracking the cells imaged via phase-contrast microscopy, Tadamoto Isogai for creating the CMV100-Arp3-Halo construct, Ugur Eskiocak, Elena Piskounova, Arin Aurora, and Sean Morrison for providing primary melanoma cells, and Bruce Kirkpatrick and Kristi Anseth for their suggestion to perform the DQ collagen experiments. We thank Stefan Wieser and Verena Ruprecht for use of their custom high NA TIRF microscope and Philippe Roudot for advice using the optical flow algorithm. We also thank Alba Diz-Muñoz and Jörg Renkawitz for kindly commenting on the manuscript. Cell and collagen rendering was performed with UCSF ChimeraX, developed by the Resource for Biocomputing, Visualization, and Informatics at the University of California, San Francisco, with support from National Institutes of Health R01-GM129325 and the Office of Cyber Infrastructure and Computational Biology, National Institute of Allergy and Infectious Diseases. This work was supported by the following grants: K25CA204526 to ESW, K99GM123221 to MKD, INSERM ITMO Cancer Plan Cancer 2015-2020 to MP, Fondation ARC no. DOC20190508743 PhD fellowship to WMG-A, NIH R33CA235254 and R35GM133522 to RF, Cancer Prevention Research Institute of Texas grant RR160057 to RF, and R35GM136428 to GD.

## Code Availability

Much of the code used in this study was associated with previously published methods papers and is available at https://github.com/DanuserLab.

---

**Fig. S1.**
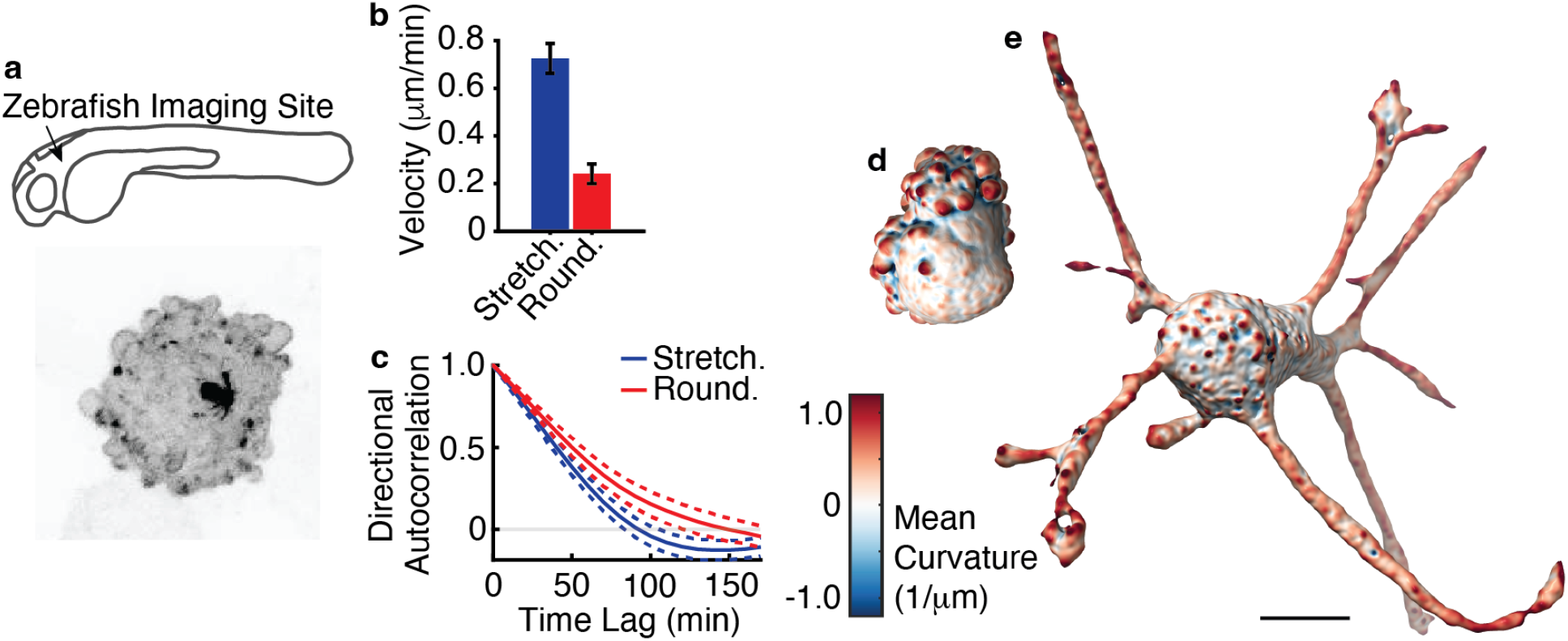
Melanoma cell morphology and migration. (a) Maximum intensity projection of a 3D confocal image of a B16 melanoma cell xenografted into a zebrafish embryo. The imaging site is indicated on the zebrafish schematic. (b) Mean instantaneous velocity of melanoma cells in 3D collagen, categorized by cell shape (n = 68 cells). (c) Directional persistence as measured by the directional autocorrelation of single cell trajectories, categorized by cell shape (n = 68 cells). Greater lag times indicate more persistent migration. Surface renderings of 3D light-sheet microscopy images of (d) amoeboid and (e) mesenchymal melanoma cells in 3D collagen. Scale bar is 10 ⍰m.

**Fig. S2.**
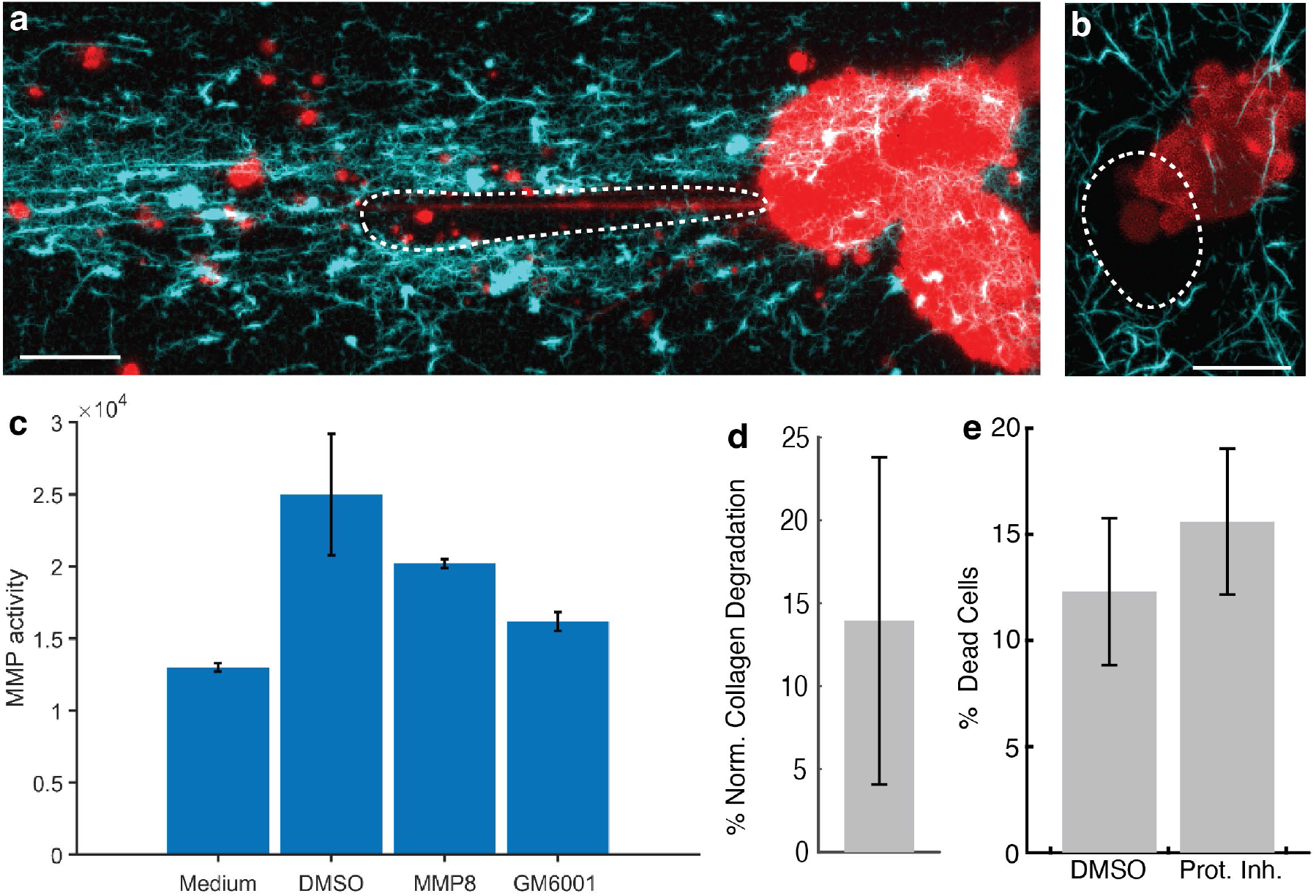
Digging tunnels through 3D collagen. (a) Maximum intensity projection across 3.2 ⍰m (20 slices) of a light-sheet microscope image of two MV3 melanoma cells expressing GFP-AktPH (red) in 3D collagen (cyan). The white dashed line indicates the location of a tunnel. (b) Maximum intensity projection across 18 ⍰m of a light-sheet microscope image of an Ewing sarcoma cell expressing GFP-F-tractin (red) in 3D collagen (cyan). The white dashed line indicates the location of a tunnel. The collagen images in a & b were computationally enhanced by a steerable line filter amplifying collinear structures, and in both panels the scale bars show 10 ⍰m. (c) MMP activity is evaluated by measuring fluorescence resonance energy transfer (FRET) fluorescence of a generic MMP peptide, measured via plate reader. In the intact FRET peptide, the fluorescence of one part is quenched by another. After cleavage into two separate fragments by MMPs, the fluorescence is recovered. ‘Medium’ shows MMP activity of DMEM cell culture medium with FBS but no cell contact. ‘DMSO’ shows activity of cell culture medium incubated with cells and DMSO vehicle as control for 24 hours. ‘MMP8’ shows the activity of purified MMP8 protein dissolved in sample measurement buffer. ‘GM6001’ shows the activity of cell culture medium incubated with cells and 40 ⍰M GM6001 for 24 hours. P value via two sample t-test for the comparison between DMSO and GM6001 is 0.0068, n=4 separate culture wells for each condition. Error bars show 95% confidence interval. (d) Collagen degradation of MV3 melanoma cells in 3D collagen treated with protease inhibitor cocktail. The degradation is normalized via a per experiment vehicle (DMSO) control corresponding to 100% collagen degradation (n=5 collagen gels). The error bar shows the standard error of the mean. (e) The percentage of visualized cells that are dead in an assay comparing protease inhibitor cocktail treated MV3 cells in 3D collagen with otherwise similar vehicle (DMSO) treated cells (n=4 collagen gels). Error bars show the standard error of the mean.

**Fig. S3.**
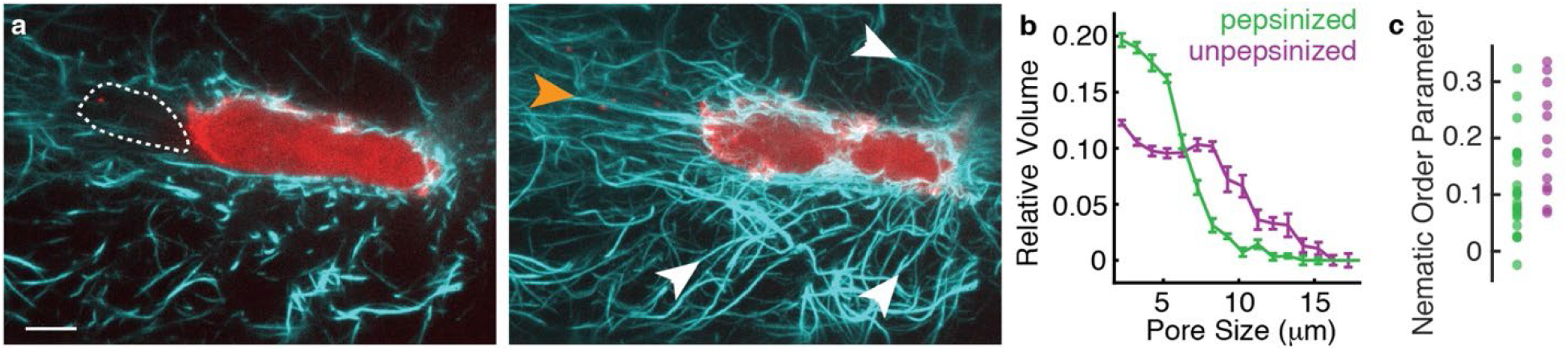
Comparing path generation in pepsinized and unpepsinized collagen. (a) Maximum intensity projections of a single light-sheet microscope image of an MV3 melanoma cell expressing GFP-AktPH (red) in pepsinized 3D collagen with fibers enhanced (cyan), projected across 1.1 ⍰m (left) and 8.2 ⍰m (right). In the left panel, the white dashed line indicates the location of a tunnel behind the cell. In the right panel, the orange arrow indicates a collagen fiber being dragged behind the cell, and the white arrows indicate collagen fibers being dragged from the sides in the cell’s wake. Scale bar show 10 ⍰m. (b) Pore size analysis of pepsinized and unpepsinized 3D collagen samples. Error bars indicate the standard error of the mean (n=6 gels per condition). The pepsinized data was shown in 1b. (c) Nematic order parameter quantifying the extent of collagen fiber alignment in images of pepsinized and unpepsinized 3D collagen containing cells (difference between conditions per a one-sided t-test, p = 0.009). A mean nematic order parameter of 1 indicates high fiber alignment and 0 indicates no overall alignment.

**Fig. S4.**
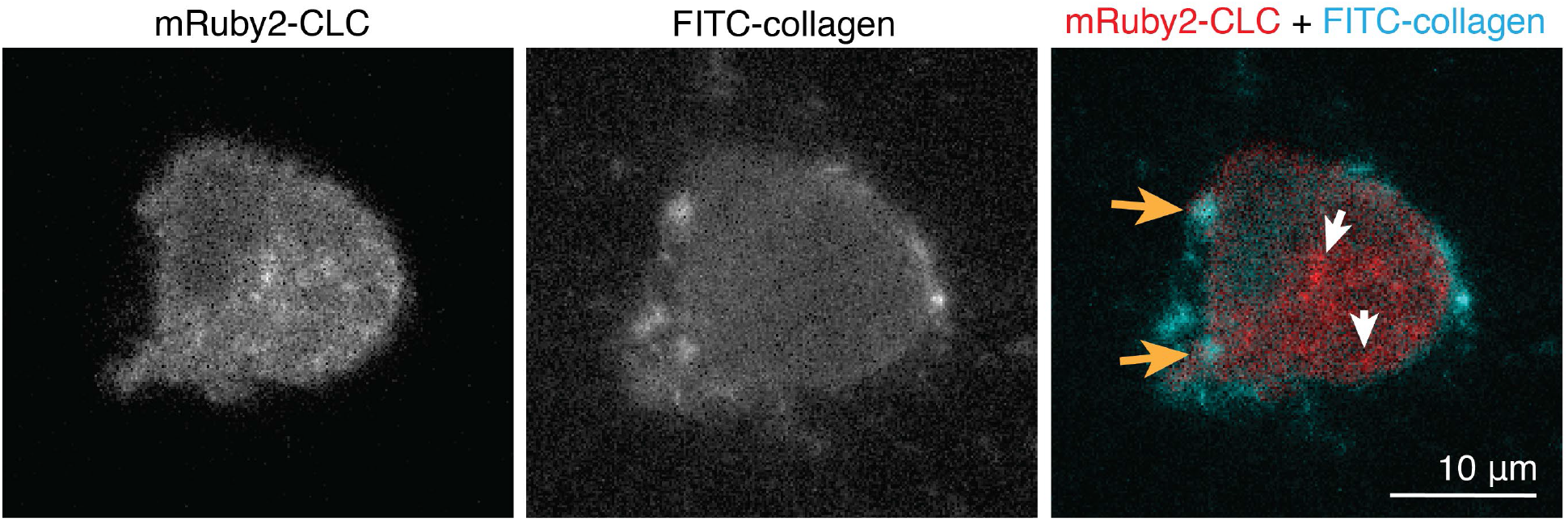
Spatial location of internalized collagen and clathrin-containing vesicles. Images show a single optical slice of a 3D light sheet microscope image of an MV3 melanoma cell expressing mRuby2-CLC in FITC-labeled 3D collagen. Orange arrows indicate internalized collagen fragments and white arrows indicate clathrin-containing vesicles.

**Fig. S5.**
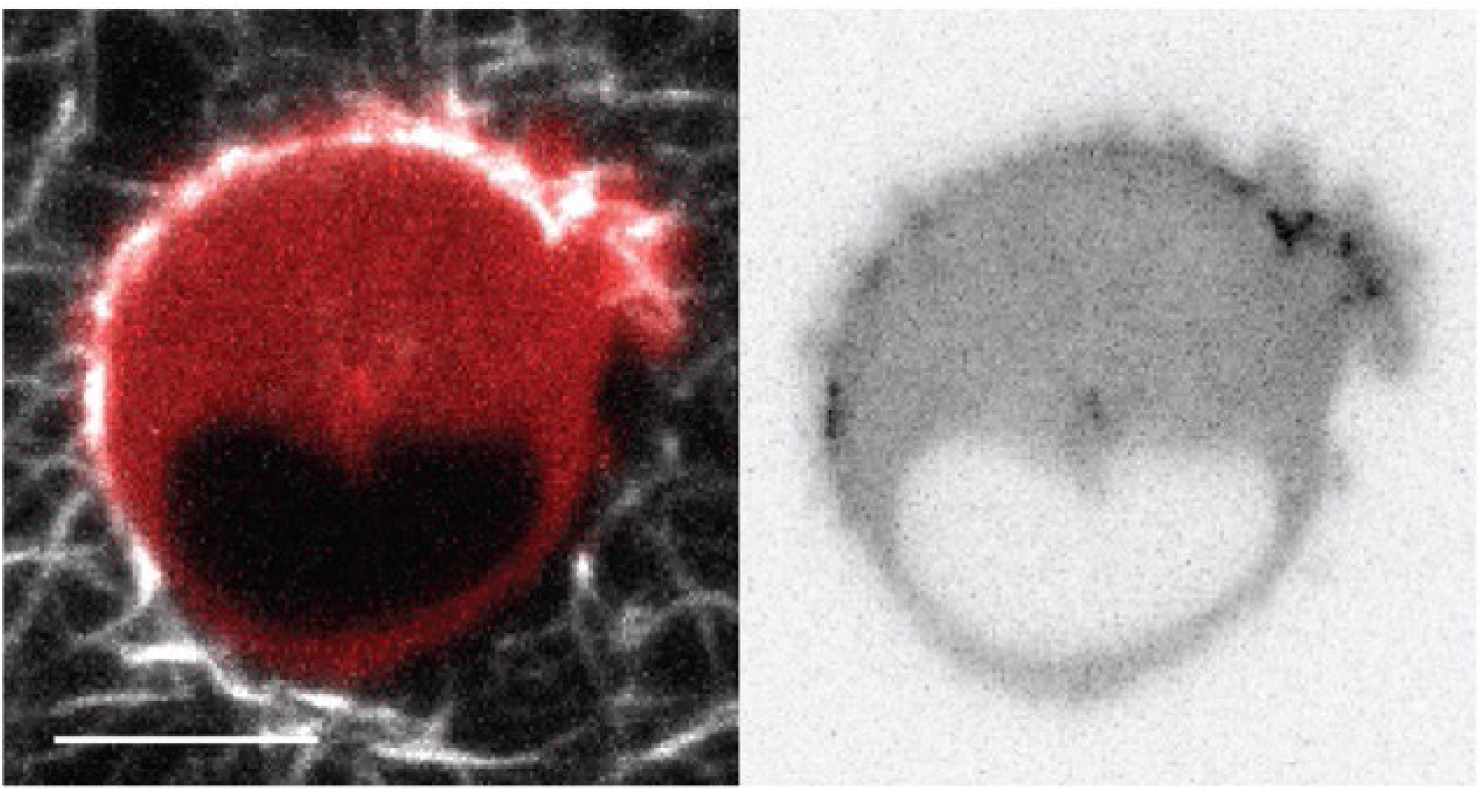
Adhesion localization. Maximum intensity projections across 0.8 ⍰m of a light-sheet microscope image of a melanoma cell expressing GFP-paxillin (red in left image, black in right image) in 3D collagen (white in left image). Scale bar indicates 10 ⍰m.

**Fig. S6.**
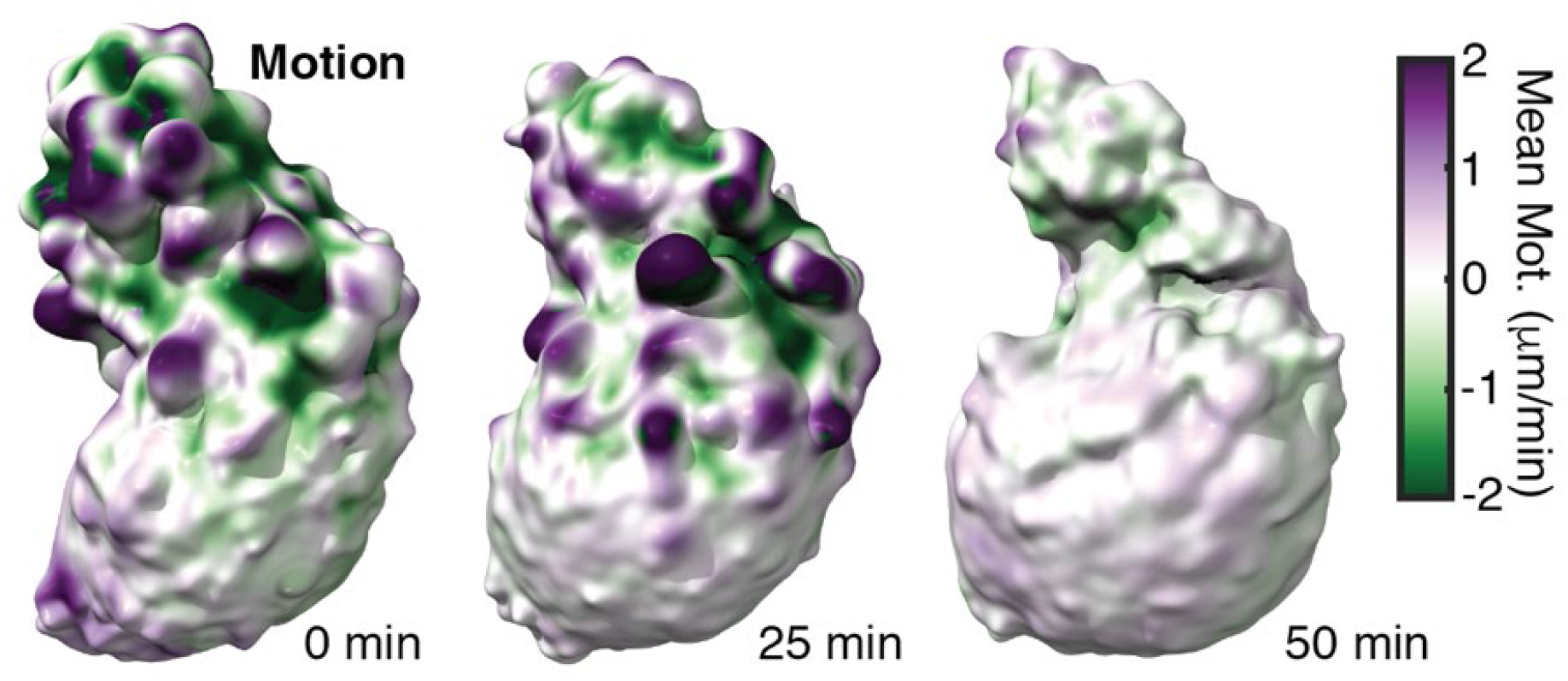
Surface motion of a melanoma cell treated with FAK inhibitor 14. Surface renderings colored by local surface motion of the cell analyzed in Figure 3K. Purple indicates protrusive regions, whereas green indicates retractive regions. Panel 3f shows PI3K activity localization on the same cell prior to treatment.

**Fig. S7.**
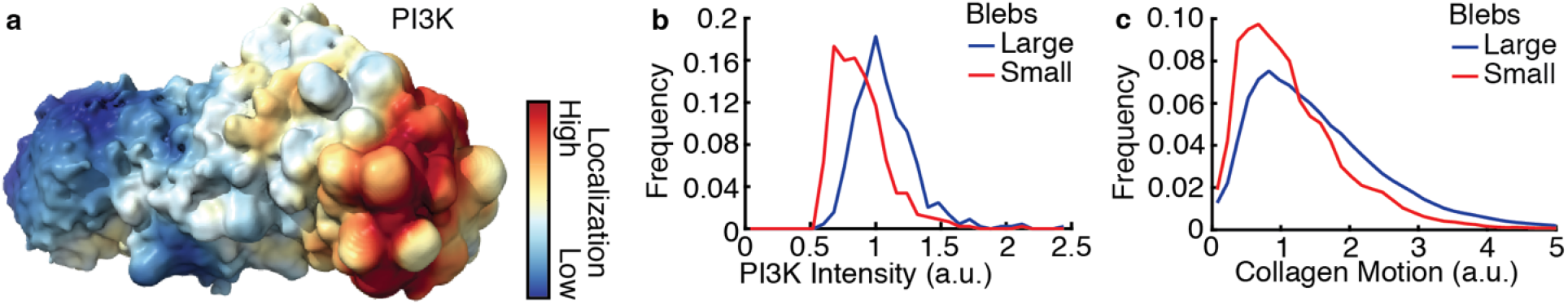
PI3K localization is associated with large blebs. (a) A surface rendering of a light-sheet microscope image of a melanoma cell in collagen, colored by the localization of GFP-AktPH. A timelapse of the same cell is shown in Movie 5. (b) Relative frequency distributions of GFP-AktPH near the cell surface reporting PI3K products in large blebs (top decile by volume) vs small blebs (bottom decile by volume) (n = 34 cells). (c) Frequency distributions of collagen motion near cell surfaces of large blebs (top decile by volume) vs small blebs (bottom decile by volume) (n = 6 cells).

**Fig. S8.**
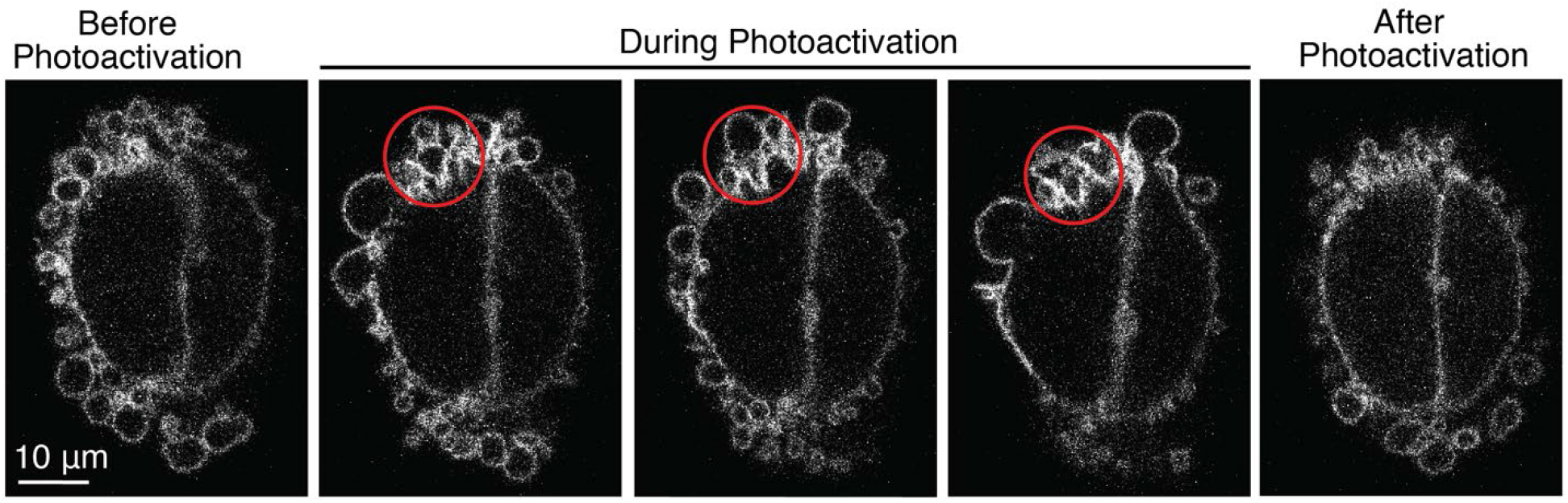
PI3K photoactivation. Spinning disk confocal microscope images showing a single optical slice of GFP-AktPH biosensor localization in an MV3 cell before, during and after photoactivation of PI3K in the area indicated by a red circle.

**Fig. S9.**
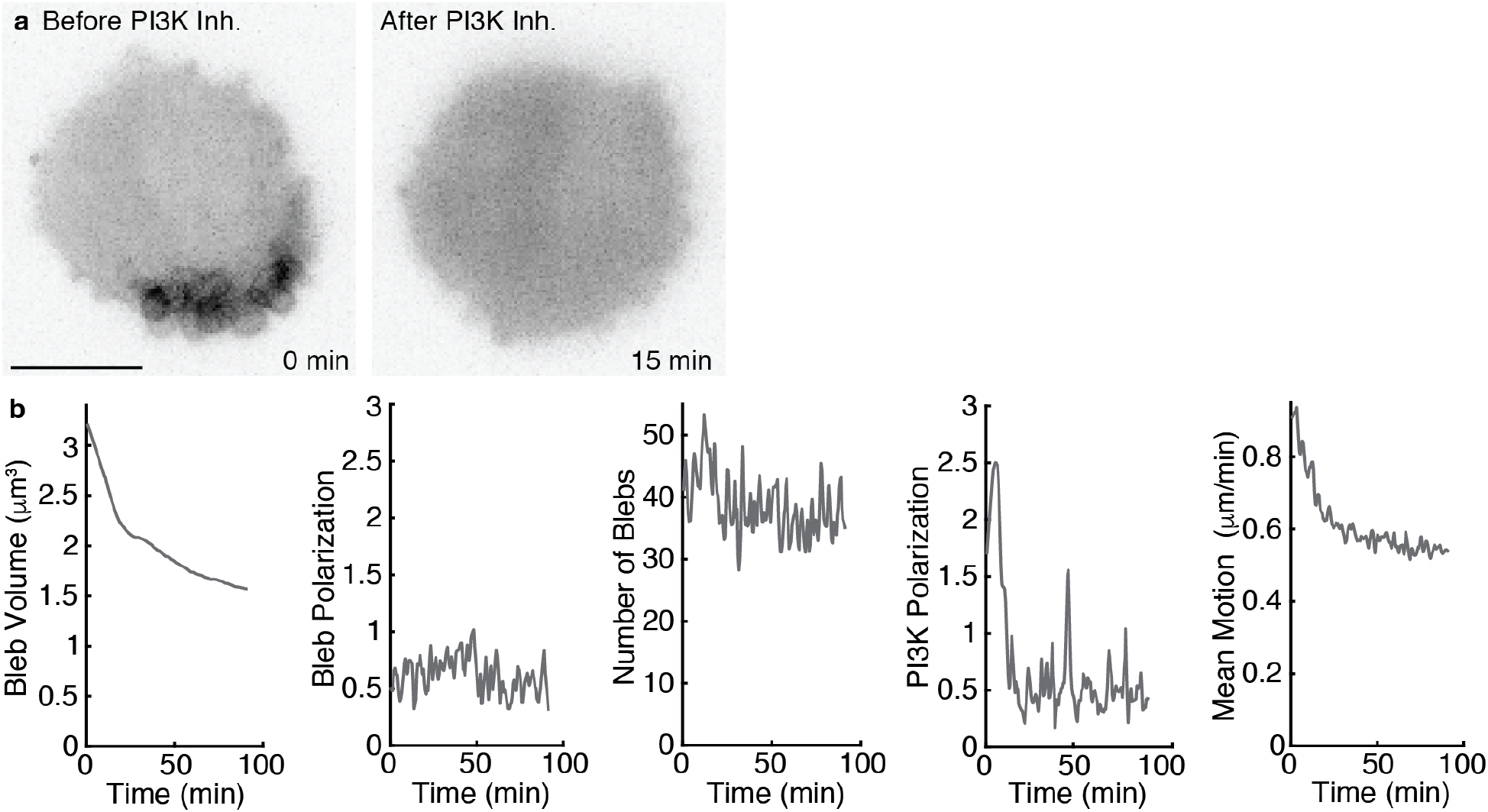
PI3K inhibition. (a) Maximum intensity projections across 1.6 ⍰m of a light-sheet microscope image of an MV3 cell expressing GFP-AktPH, before and after PI3K inhibition. Scale bar shows 10 ⍰m. (b) Temporal response of a different MV3 cell treated with PI3K⍰ inhibitor IV. From left to right, shown are mean bleb volume, bleb polarization magnitude, number of blebs, PI3K polarization magnitude, and mean surface motion magnitude.

**Fig. S10.**
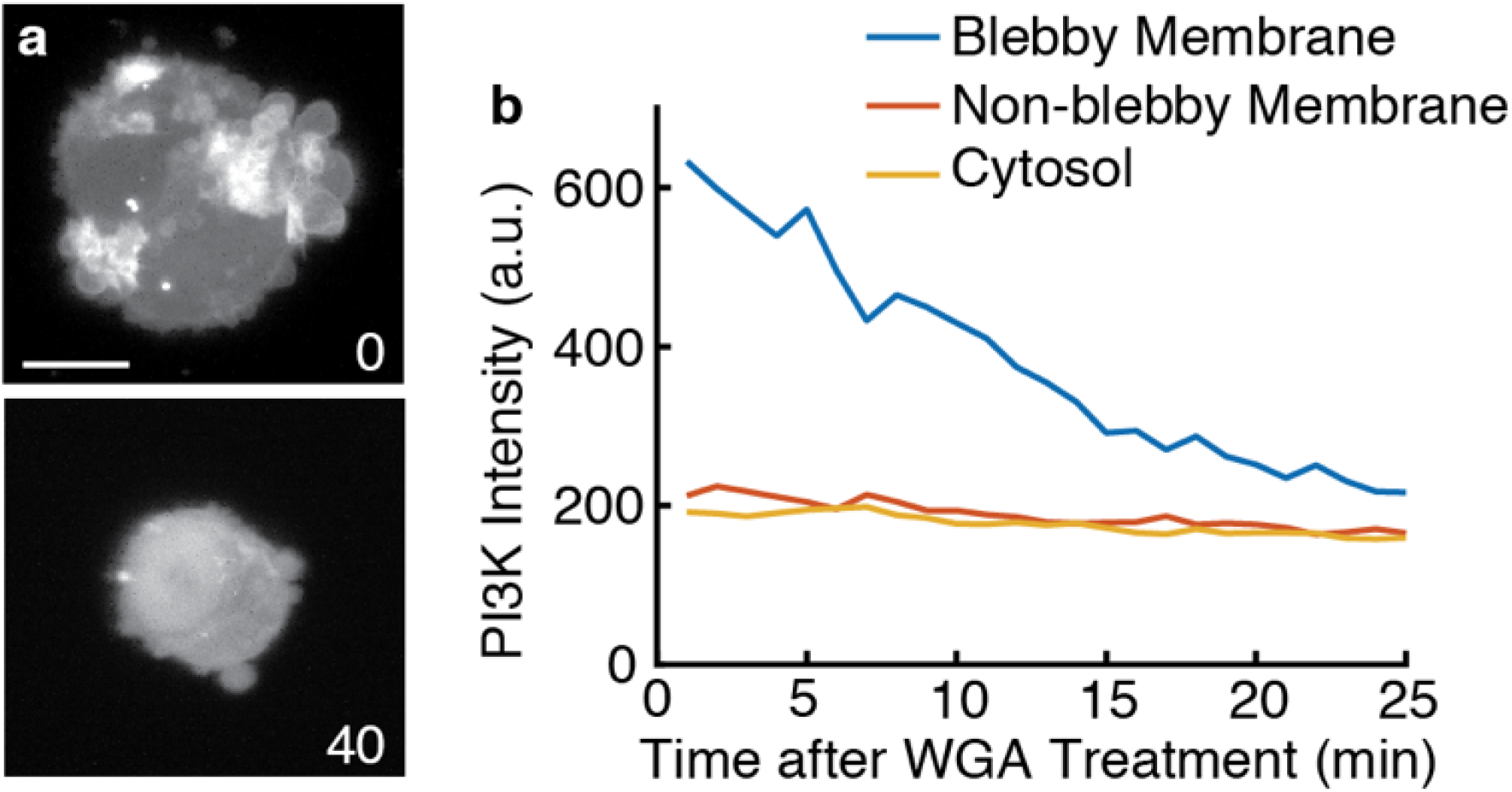
Wheat Germ Agglutinin (WGA) treatment. (a) Maximum intensity projections of light-sheet microscope images of representative melanoma cells treated with 0 ⍰g/mL or 40 ⍰g/mL WGA. Scale bar shows 10 ⍰m. (b) PI3K intensity as a function of time after WGA treatment in different regions of a cell.

**Fig. S11.**
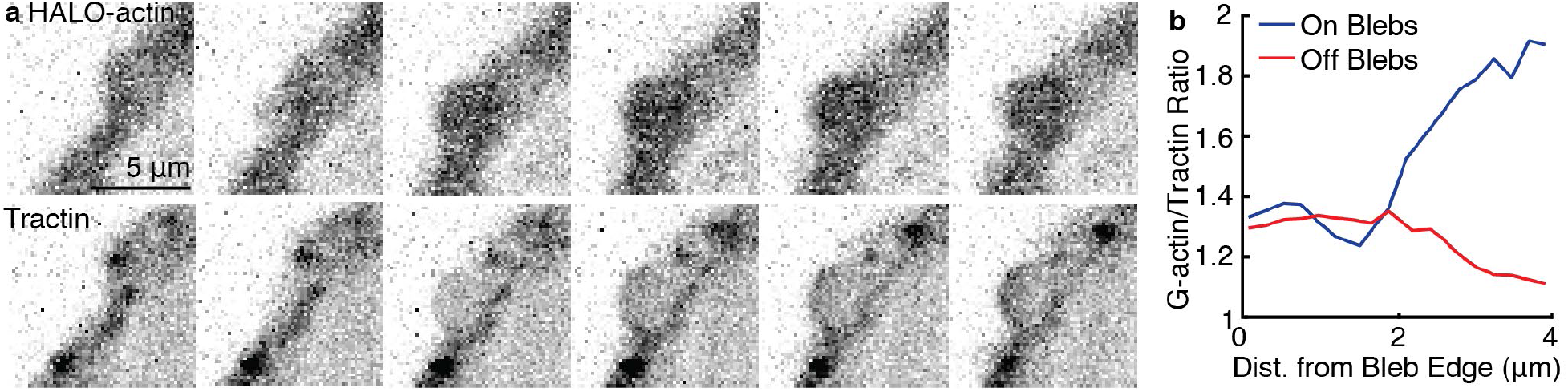
HALO-actin and tractin localization within blebs. (a) Time-lapse maximum intensity projections across 3.2 ⍰m of HALO-actin (total actin) and GFP-F-tractin in an MV3 cell. Volumes were acquired every 4.3 sec. (b) Actin/tractin ratio on and off blebs as a function of distance from the bleb neck (n = 5 cells).

**Movie S1. Collagen movement in soft collagen.** Single optical slice of a light sheet fluorescence time-lapse series showing motion of collagen labeled with Alexa Fluor 568 in the proximity of a single melanoma cell (unlabeled).

**Movie S2. Collagen network surrounding a melanoma cell.** Movie animates stepping through single slices of a light sheet fluorescence image of collagen labeled with Alexa Fluor 568 along with a single melanoma cell showing local surface curvature mapped to the surface of the cell.

**Movie S3. A cell tunneling through collagen.** Movie shows a time-lapse sequence of a single slice of a light sheet microscope image of a melanoma cell expressing GFP-AktPH (green) and collagen (red).

**Movie S4. Animation showing blebs identified by u-shape 3D.** Movie shows rotation of a 3D rendering of the surface of a single melanoma cell, imaged using light sheet microscopy. Blebs identified by u-shape 3D are shown in different colors.

**Movie S5. PI3K polarity in a migrating cell.** Movie shows a time-lapse sequence of the surface of light sheet microscope images of a melanoma cell expressing GFP-AktPH. The local intensity of GFP-AktPH is projected onto the surface of the cell, according to the colormap shown in Extended Data Figure 7a.

**Movie S6. Photoactivation of PI3K in a blebbing cell.** Time-lapse sequence of a single optical section of a confocal microscope acquisition of an MV3 cell expressing GFP-AktPH. Photoactivation of Cry2-iSH2/CIBN-CAAX occurred via activation of the FRAP module in the location and time shown by the green circle.

**Movie S7. PI3K inhibition in a melanoma cell with polarized PI3K and blebs.** Movie shows a time-lapse sequence of a maximum intensity projection of light sheet microscope images of a melanoma cell expressing GFP-AktPH. Images are inverted so that dark areas represent high PI3K activity. 500 nM of PI3K inhibitor was added at time 0.

**Movie S8. Time-lapse sequence of a single bleb in a HeLa cell expressing GFP-actin.** Images were acquired via TIRF microscopy.

**Movie S9. Photoactivation of Rac1 in a blebbing cell.** Time-lapse sequence of a single optical section of a confocal microscope acquisition of a mouse embryonic fibroblast cell expressing mCherry-PARac1. Photoactivation occurred in the location and time shown by the red circle.

**Movie S10.** Automatic bleb tracking during Tiam1 photoactivation in melanoma cells. The MV3 cells are expressing sspb tagRFP Tiam1, with the dashed line indicating the region and timing of photoactivation. This movie is 10 minutes long (200 frames).

## Notes

### Competing Interest Statement

The authors have declared no competing interest.

### Summary of Updates

Additional data on the independence of the worrying migration form of matrix metalloprotease activity. Additional optogenetic experiments to support the Rac-signaling dependence of bleb expansion.

